# Improved detection of tumor suppressor events in single-cell RNA-Seq data

**DOI:** 10.1101/2020.07.04.187781

**Authors:** Andrew E. Teschendorff, Ning Wang

## Abstract

Tissue-specific transcription factors are frequently inactivated in cancer. To fully dissect the heterogeneity of such tumor suppressor events requires single-cell resolution, yet this is challenging because of the high dropout rate. Here we propose a simple yet effective computational strategy called SCIRA to infer regulatory activity of tissue-specific transcription factors at single-cell resolution and use this tool to identify tumor suppressor events in single-cell RNA-Seq cancer studies. We demonstrate that tissue-specific transcription factors are preferentially inactivated in the corresponding cancer cells, suggesting that these are driver events. For many known or suspected tumor suppressors, SCIRA predicts inactivation in single cancer cells where differential expression does not, indicating that SCIRA improves the sensitivity to detect changes in regulatory activity. We identify NKX2-1 and TBX4 inactivation as early tumor suppressor events in normal non-ciliated lung epithelial cells from smokers. In summary, SCIRA can help chart the heterogeneity of tumor suppressor events at single-cell resolution.

## Introduction

Tissue-specific transcription factors are required for the differentiated state of cells in a given tissue ^1^. They are often inactivated in cancer, which is associated with a lack of differentiation, a well-known cancer hallmark ^2–6^. Many of these tissue-specific transcription factors (TFs) encode tumor suppressors and their inactivation may constitute driver events that are thought to occur in the earliest stages of carcinogenesis ^7–9^. Estimating regulatory activity of such tissue-specific transcription factors (TFs) in both normal and cancer tissue is therefore a critically important task, as this can reveal which normal tissues are at risk of neoplastic transformation ^10^. There are two main reasons why this task should be performed at single-cell resolution ^11–13^. First, TFs control cell-identity ^1,14^, and thus, estimation of regulatory activity in bulk tissue is subject to confounding by cell-type heterogeneity. Second, to fully characterize cancer heterogeneity requires identifying putative tumor suppressor events at the most fundamental scale, i.e. the single-cell ^15–18^.

However, estimating regulatory activity of TFs at single-cell resolution is hard, because of the typically high dropout rate and low genomic coverage of single-cell assays ^19–21^. In the context of single-cell RNA-Seq assays, one could in principle use TF expression as a surrogate marker of TF-activity (i.e. regulatory activity reflecting the effect of the TF on downstream expression of direct and indirect targets), and while this strategy works well on expression data derived from bulk tissue (see e.g. ^1^), it is unclear how well this works for scRNA-Seq assays ^22,23^. Thus, it is also unclear how best to infer regulatory activity in the majority of scRNA-Seq cancer studies that are performed in solid epithelial tissues.

Here we present a novel strategy called SCIRA (SCalable Inference of Regulatory Activityin single cells), which applies an existing regulatory inference method ^8^ to a suitably powered bulk multi-tissue RNA-Seq dataset to identify tissue-specific TFs and their regulons (i.e. their direct and indirect targets), from which regulatory activity in single cells can then be estimated. We comprehensively validate SCIRA and demonstrate through a power calculation and application to real scRNA-Seq data, that SCIRA can estimate regulatory activity even for TFs that are highly expressed only in relatively minor fractions (~5%) of cells within a bulk tissue. We subsequently apply SCIRA to several scRNA-Seq datasets containing both normal and cancer cells, where it reveals preferential inactivation of tissue-specific TFs in corresponding single cancer cells, an observation strongly consistent with analogous results obtained in bulk tissue ^5^, whilst also revealing novel tumor suppressor events at single-cell resolution. We further showcase an important application of SCIRA to identify tumor suppressor events in single normal cells (lung epithelial cells) exposed to a cancer risk factor (smoking). Our results underscore the critical need for a method like SCIRA, since ordinary differential expression fails to reveal the same insights, even after imputation of dropouts.

## Results

### Inferring regulatory activity with SCIRA: rationale

SCIRA identifies tissue-specific TFs, builds regulons for these TFs, and uses these regulons to estimate regulatory activity of the TFs in scRNA-Seq data (**Methods**). SCIRA adapts the SEPIRA algorithm (previously published by us ^8^) to infer tissue-specific TFs and regulons from the large GTEX multi-tissue bulk RNA-Seq dataset (8555 samples, 30 tissue-types) ^24^ (**Methods, Fig.1A**). We note that the tissue-specific TFs are derived by adjusting for cell-type (stromal) heterogeneity, which can otherwise strongly confound differential expression analyses (**Methods**) ^25^. To justify inferring TFs and their regulons from bulk tissue data, we performed a careful power calculation, which revealed that SCIRA has reasonable sensitivity to detect tissue-specific TFs that are highly expressed even if only in a relatively underrepresented cell-type within the tissue (**Methods, Fig.1B**). For instance, using reasonable values for the average fold-change (**SI fig.S1**), we estimated that for tissues like lung, pancreas and liver, for which there are more than 100 samples in GTEX (total number of samples is 8555), sensitivity to detect TFs expressed in only 5% of cells within the tissue (i.e. a minor cell fraction MCF=0.05) were generally still over 50% (**Fig.1B**, **SI fig.S2**). The inferred TF-regulons can subsequently be applied to suitably-matched scRNA-Seq data in a linear regression framework ^26^ (**Methods**) to estimate regulatory activity for each single cell. By using the actual regulon of the TF, this inference should be robust to dropouts, i.e. even if the TF itself is not detected across most if not all of the cells in the study (**Fig.1C**). Finally, one can construct regulatory activity maps across the relevant cells within the tissue (**Fig.1C**), which can reveal deregulated TFs at single-cell resolution.

**Figure-1:**
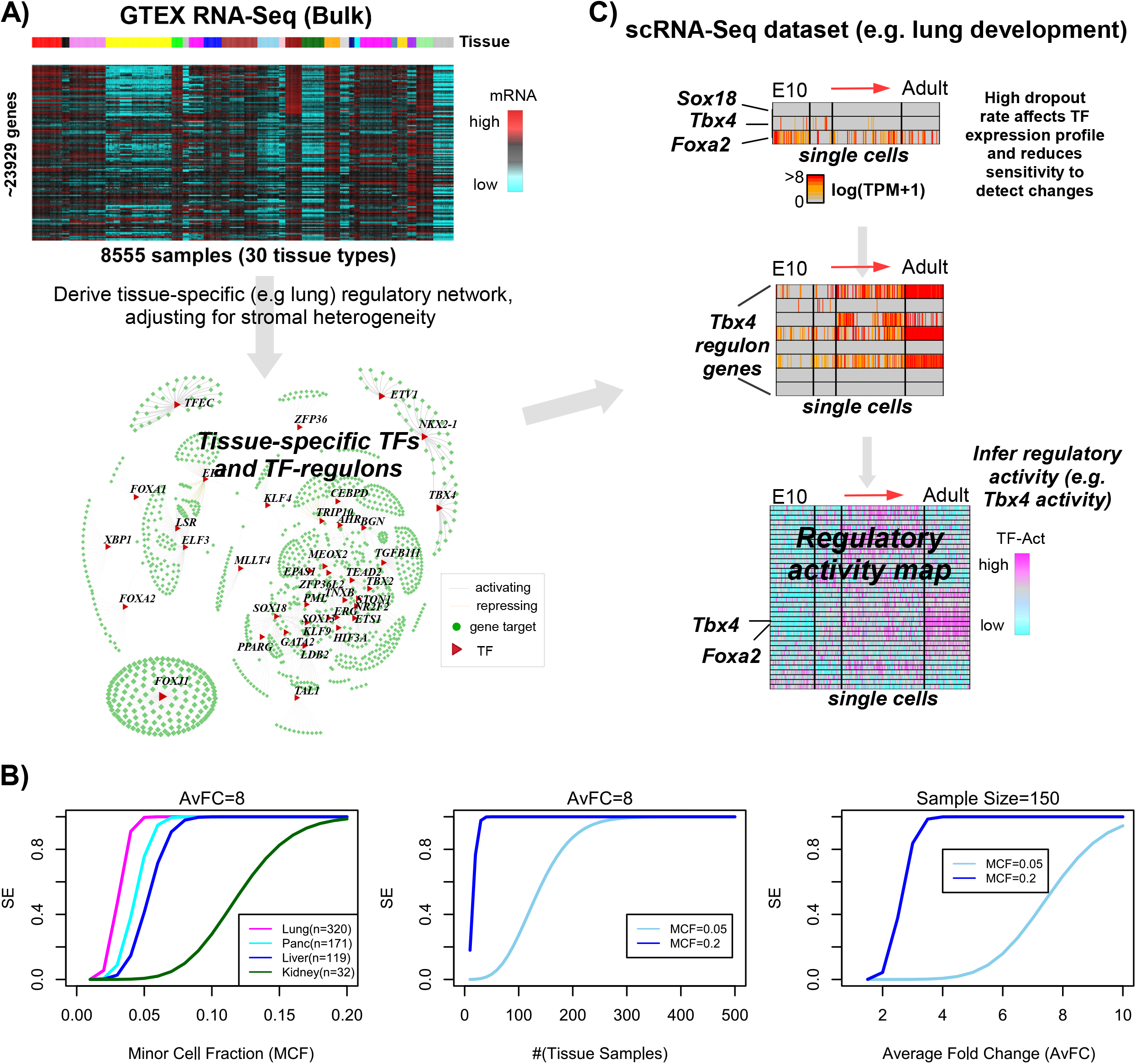
SCIRA rationale and workflow. **A)** Since bulk RNA-Seq data does not suffer from technical dropouts and is much more reliable than scRNA-Seq data, for a given choice of tissue, we use the high-powered GTEX bulk RNA-Seq expression set (>20,000 genes, 8555 samples, 30 tissue-types) to derive a corresponding tissue-specific regulatory network, consisting of a gold-standard list of tissue-specific transcription factors (TFs) and their targets (regulons). The inference of the network uses a greedy partial correlation framework, whilst also adjusting for stromal (immune cell) contamination within the tissue. **B)** Power/Sensitivity (SE) estimates to detect tissue-specific TFs in the GTEX bulk RNA-Seq dataset as a function of the minor cell-type fraction (MCF) (left), number of samples in the tissue of interest (middle) and average fold change of differential expression between the tissue of interest and the rest of tissues in GTEX (right). In left panel, we depict SE curves for 4 tissue types in GTEX (number of samples in each tissue is given) and for an average FC=8. In the middle panel we depict SE curves for two MCF values, as indicated. In the right panel, we assume a sample size of 150. A MCF value of 0.05 means we assume that the tissue-specific TFs is only highly expressed in 5% of the tissue resident cells. **C)** Given the high technical dropout rate and overall noisy nature of scRNA-Seq data, it may not be possible to reliably infer regulatory activity from the TF expression profile alone. However, using the TF regulons derived in A), and using the genes within the regulon that are not strongly affected by dropouts, we can estimate regulatory activity across single-cells. Depicted is an example with 3 lung-specific TFs (*Sox18, Tbx4*, *Foxa2*), as well as the expression pattern of the regulon genes for *Tbx4,* in the context of a lung development study from embyronic day-10 to adult stage (Treutlein dataset). We use linear regressions between the expression values of all the genes in a given cell and the corresponding TF-regulon profile, to derive the activity of the TF as the t-statistic of the estimated regression coefficient, resulting in a regulatory activity map over the tissue-specific TFs and single cells. The same tissue-specific TFs and their regulons can be applied to normal-cancer scRNA-Seq datasets to infer regulatory activity maps across normal and cancer cells.

### Validation of SCIRA in normal tissue

As a proof of principle we applied SCIRA to four tissue-types (lung, liver, kidney and pancreas) using the GTEX dataset to infer corresponding tissue-specific TFs and regulons. We identified on average about 30 tissue-specific TFs for each of the 4 tissue-types and on average about 40 to 50 regulon genes per TF (**SI tables S1-S4, Supplementary File 1).** The TF lists contained well-known tissue-specific factors: e.g. for liver, the list included the well-known hepatocyte factors *HNF1A, HNF4A* and *FOXA1 (HNF3A);* for lung, the list included well-known lung alveolar differentiation factors *TBX2* and *FOXA2* ^27–29^, and *FOXJ1*, a factor required for ciliogenesis ^30^. In order to test the reliability of the TFs and regulons, we performed four separate validation analyses.

First, although there is no logical requirement for regulon genes to be direct targets ^31^, some enrichment for direct binding targets is expected. Approximately 65% of our TF-regulons exhibited statistically significant enrichment for corresponding ChIP-Seq TF-binding targets (**SI fig.S3-S4**), as determined using data from the ChIP-Seq Atlas ^32^ (**Methods**). For instance, in the case of liver we could find ChIP-Seq data for 12 of the 22 liver-specific TFs, and for 9/12 we observed statistically significant enrichment (**SI fig.S3D-E**). In many instances, proportions of regulon genes that were direct TF binding targets were considerable. For example, for the liver-specific TF *HNF4G*, 57% of its 37 regulon genes (i.e. 21 genes) were bound by *HNF4G* within +/− 5kb of the gene’s transcription start site (TSS) (**SI fig.S3D**). For *FOXA1,* 8 of its 10 regulon genes were bound by *FOXA1* within +/−1kb of the TSS (**SI fig.S3D**). Statistical significance estimates were independent of the choice of threshold on binding intensity values (**Methods**), and also robust to parameter choices in SCIRA (**SI fig.S5, Methods**). Second, we were able to validate the tissue-specificity of the regulons and derived regulatory activity estimates in independent multi-tissue bulk RNA-Seq (ProteinAtlas ^33^) and microarray data from Roth et al ^34^ (**SI fig.S6-S9**). Given these successful validations, we estimated on average only 10% of TF regulon-gene associations to be false positives (**SI fig.S10**). Third, we collated and analysed scRNA-Seq datasets representing differentiation timecourses into mature epithelial cell-types present within the given tissues, encompassing two species (human & mouse) and 3 different single-cell technologies (Fluidigm C1, DropSeq & Smart-Seq2) (**SI table S5, Methods**) ^35–38^. We reasoned that most of our tissue-specific TFs would exhibit higher regulatory activity in the corresponding mature differentiated cells compared to the immature progenitors, a hypothesis that we were able to strongly validate in each of the four tissue-types (**SI fig.S11-S14**). These results could not have arisen by random chance and were not seen if we used tissue-specific TFs from other unrelated (non-epithelial) tissues like skin or brain (**SI fig.S15**). We further observed that, owing to the high dropout rate, SCIRA’s regulatory activity estimates were much more sensitive than expression itself (**SI fig.S11-S14, Fig.2A**). As a concrete example, SCIRA’s regulatory activity estimates for lung alveolar differentiation factors *TBX2* and *FOXA2* ^27–29^ were higher in the mature alveolar cell-types compared to the immature progenitors, as required, whilst expression levels could not detect an increase (**SI fig.S11**). SCIRA displayed improved sensitivity and prevision (i.e. lower false discovery rate) over differential expression (DE) even after application of imputation methods (scImpute ^39^, MAGIC ^40^, Scrabble ^41^), or even when compared to other regulatory activity estimation methods like SCENIC/GENIE3 ^42^ (**Fig.2A-C, Methods**). SCIRA also displayed improved sensitivity over the combined use of VIPER ^43,44^ and the dorothea TF-regulon database ^45,46^ (“VIPER-D”), as well as lower FDRs (**Fig.2A-C, Methods**). This is noteworthy given that the TF-regulons from dorothea are not tissue-specific. Fourth, we validated the power calculation underlying SCIRA by applying it to a differentiation timecourse scRNA-Seq dataset in liver ^36^, which revealed the expected bifurcation of hepatoblasts into hepatocytes and cholangiocytes, as well as identifying cholangiocyte specific factors, despite their very low frequency (5-10%) in liver tissue (**SI fig.S16B-E, SI fig.S17).** We note that the bifurcation and dynamic expression patterns were not revealed when analyzing TF expression levels (**SI fig.S18**), further supporting the view that SCIRA can improve the sensitivity to detect correct patterns of TF-activity.

**Figure-2:**
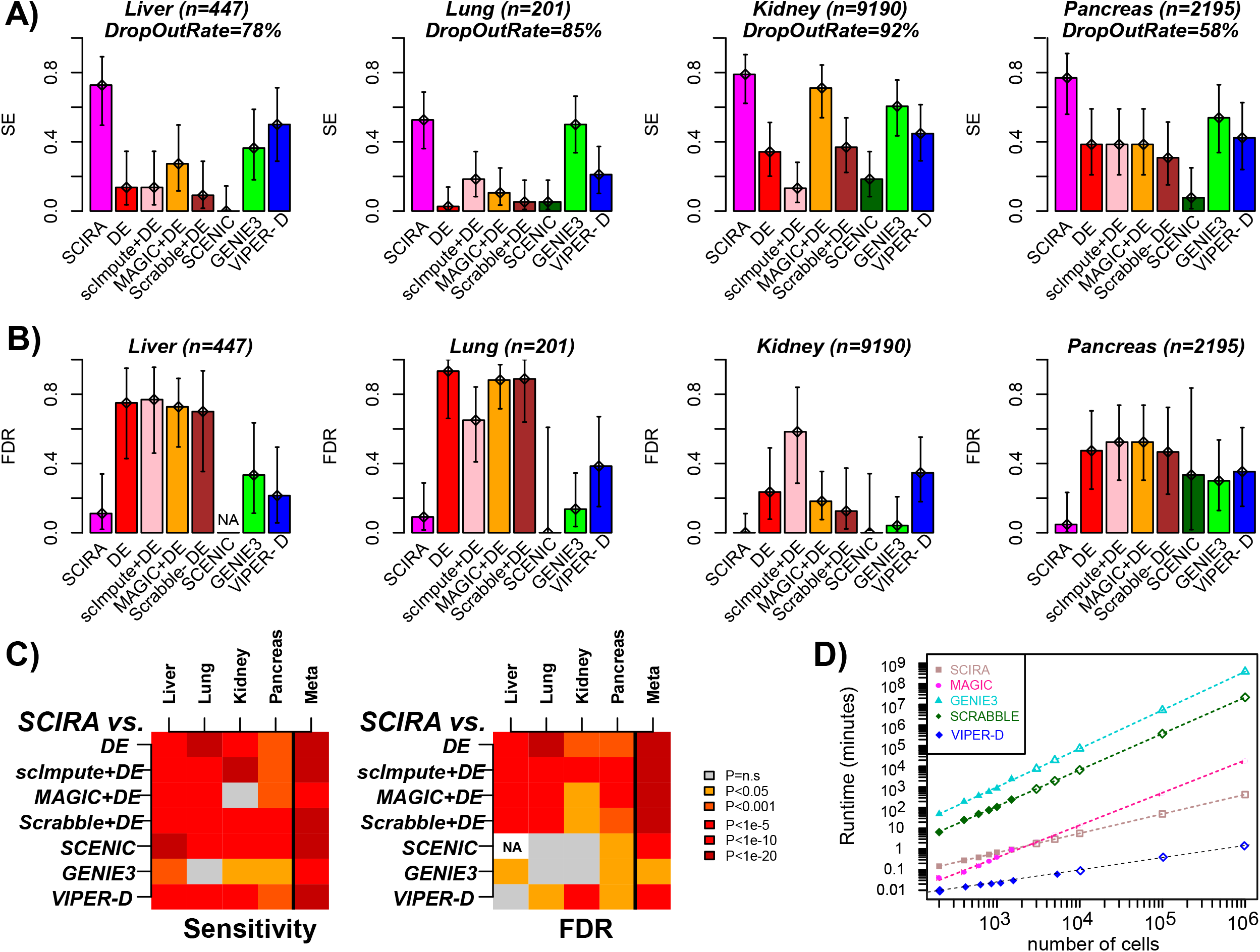
SCIRA displays improved sensitivity, precision and scalability. **A)** Barplots with 95% confidence intervals included displaying the sensitivity (SE) to detect increased activity or expression for a gold-standard set of tissue-specific TFs in a corresponding timecourse differentiation scRNA-Seq study. Methods represented are SCIRA, ordinary differential expression (DE), imputation with scImpute, MAGIC or Scrabble following by DE, SCENIC, running SCENIC without the TF-binding motif enrichment step (denoted “GENIE3”) and VIPER using the dorothea regulon database (denoted “VIPER-D”). **B)** Barplots and 95% confidence intervals displaying the false discovery rate (FDR) of each method in the same scRNA-Seq datasets. Precision is defined as 1-FDR and is the fraction of true positives among all positives. In this case, tissue-specific TFs predicted to be significantly downregulated/inactivated during the timecourse were identified as false positives with FDR defined as the fraction of false positives among all significantly differentially expressed (or activated) TFs. **C)** Heatmap of P-values assessing the improvement of SCIRA over the other 7 methods, in terms of both sensitivity (left) and FDR (right). P-values for each tissue were derived from a one-tailed Binomial test. The P-values for the meta-analysis (“Meta”) were derived using Fisher’s method. The FDR for SCENIC in liver could not be defined as the number of positives was zero. **D)** A plot of run times (y-axis, log-scale) for 5 methods (SCIRA, MAGIC, Scrabble, GENIE3/SCENIC and VIPER-D) against the number of single-cells profiled (x-axis, log-scale). Filled symbols represent times estimated from actual runs, unfilled symbols are imputed estimates obtained by extrapolation of fitted linear functions (on a log-scale). Run times were estimated using 4 processing cores (SCIRA, MAGIC, GENIE3/SCENIC, VIPER-D) and 1 core for Scrabble (as Scrabble offers no option for parallelization).

### SCIRA predicts inactivation of tissue-specific TFs in corresponding tumor epithelial cells

Next, we applied SCIRA to a recent lung cancer scRNA-Seq study (Lambrecht et al) ^47^ which profiled a total of 52,698 cells (10X Chromium) derived from 5 lung cancer patients (2 lung adenoma carcinomas – LUAD, 2 lung squamous cell carcinomas – LUSC and 1 non-small cell lung cancer-NSCLC). We hypothesized that many of our previously identified lung-specific TFs would be inactivated in lung epithelial tumor cells ^5,8^, since lack of differentiation is a well-known cancer hallmark ^6^. We used the same dimensional reduction and tSNE-approach as in Lambrecht et al ^47^, to first categorize specific clusters of cells as normal alveolar epithelial (n=1709) and tumor epithelial (n=7450) (**Fig.3A**). We verified that the alveolar cells expressed relatively high levels of an alveolar marker (*CLDN18*) (**Fig.3B**), whilst both alveolar and tumor epithelial cells expressed relatively high levels of *EPCAM*, a well-known epithelial marker (**Fig.3C**). As noted by Lambrecht et al, the great majority of alveolar cells were from non-malignant specimens representing normal (squamous) epithelium and clustered together irrespective of patient-ID ^47^, whilst cancer cells clustered according to patient (**Fig.3A**) ^47^. Next, we used SCIRA to estimate regulatory activity for all 38 lung-specific TFs in each of the (1709+7450) cells, and computed t-statistics of differential activity between alveolar and tumor epithelial cells. Remarkably, 35 out of the 38 TFs exhibited a statistically significant (Bonferroni adjusted P < 0.05) reduction in regulatory activity in tumor cells (**Fig.3D**, Wilcox-test P<1e-8). Using 1000 Monte-Carlo randomizations of the regulons, we verified that this number of inactivated TFs could not have arisen by chance (**Fig.3D,** Monte Carlo P<0.001). Among the most significantly inactivated TFs, we observed *FOXA2*, a TF required for alveolarization and which regulates airway epithelial cell differentiation ^28,29^ (**Fig.3E**), and *NKX2-1*, a master TF of early lung development ^48^ (**SI fig.S19)**. Other inactivated TFs included *SOX13,* which has been broadly implicated in lung morphogenesis ^49^, *HIF3A*, which has been shown to be highly expressed in alveolar epithelial cells and thought to be protective of hypoxia-induced damage ^50^, and the aryl hydrocarbon receptor (*AHR*), which is a regulator of mucosal barrier function and activation of which enhances CD4+ T-cell responses to viral infections ^51,52^ (**SI fig.S19**). Importantly, these findings would not have been obtained had we performed DE or VIPER-D analysis on the 38 TFs (**Fig.3D & 3F**). Indeed, according to a Wilcoxon rank sum test, 21 TFs were differentially expressed between alveolar and tumor epithelial cells, but with no clear trend towards underexpression in tumor cells (**Fig.3D**). For instance, according to single-cell DE analysis, TFs such as *TBX4* and *FOXJ1*, both with important roles in lung tissue development, were not underexpressed in tumor cells, yet found to be inactivated according to SCIRA (**Fig.3F**). Given that *TBX4* and *FOXJ1* have been found to be inactivated/underexpressed in bulk lung cancer tissue ^8^, this further supports the view that SCIRA improves sensitivity over ordinary DE analysis. To explore this further we compared the differential activity and differential expression patterns between normal and cancer cells to the differential expression patterns in the two TCGA lung cancer studies ^53,54^. A stronger agreement with the bulk RNA-Seq data of both TCGA cohorts was observed for SCIRA’s differential activity profiles compared to differential expression or when using VIPER-D to infer differential activity (**Fig.3F-G**). Indeed, approximately 30 of the 38 TFs exhibited differential activity patterns at the single-cell level that were consistent with differential expression in bulk, whilst for differential expression and VIPER-D this number was only around 10 (**Fig.3H**).

**Figure-3:**
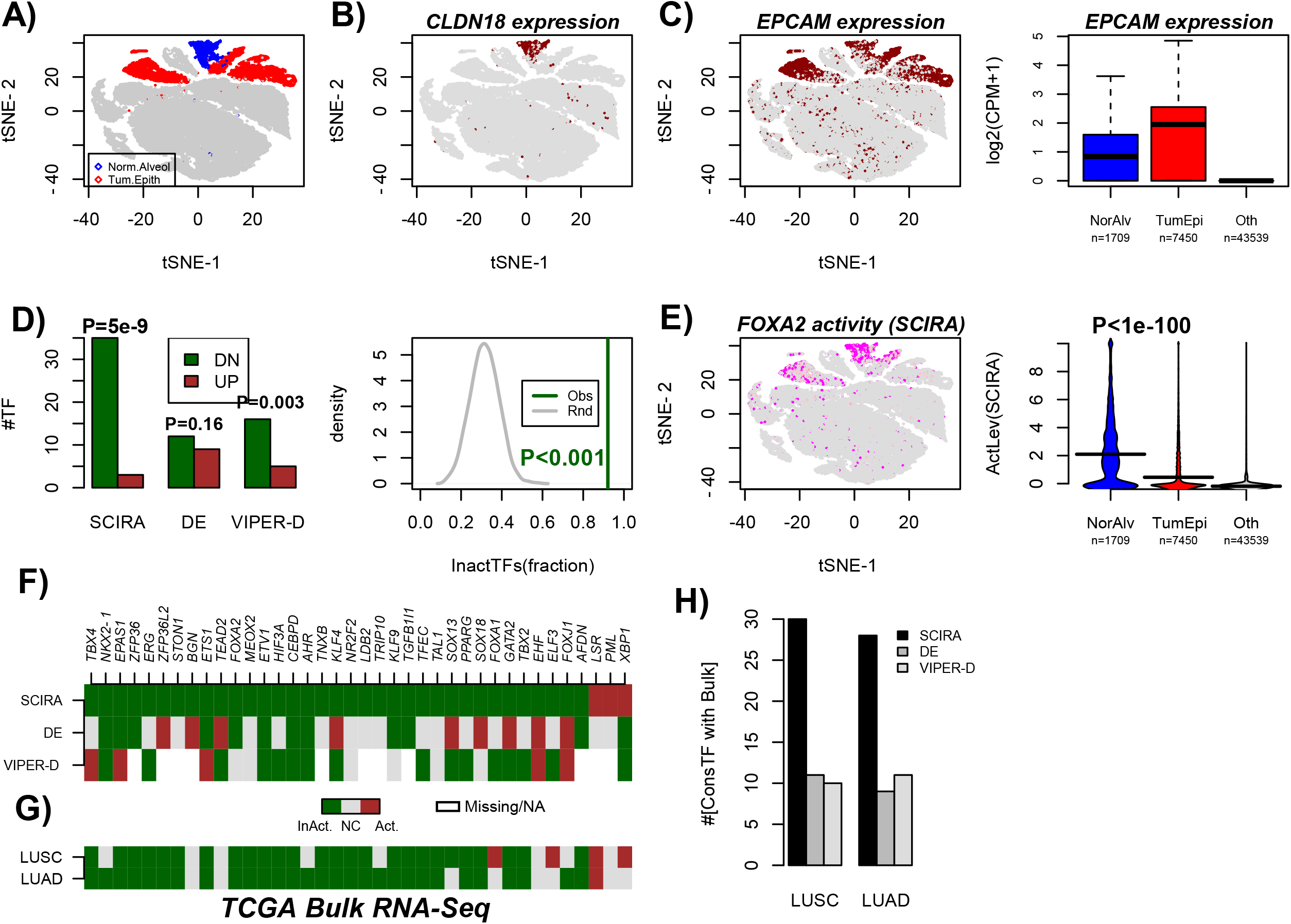
SCIRA predicts inactivation of lung-specific TFs in lung tumor epithelial cells. **A)** t-SNE scatterplot of approximately 52,000 single cells from 5 lung cancer patients, with a common non-malignant alveolar and (tumor) epithelial clusters highlighted in blue and red, respectively. **B)** Corresponding t-SNE scatterplot with cells colored-labeled by expression of an alveolar marker *CLDN18.* **C)** As B), but with cells colored according to expression of the epithelial marker *EPCAM.* Right panel depicts boxplots of the log_2_(counts per million + 1) of *EPCAM* for cells in the non-malignant alveolar cluster, the tumor epithelial clusters and all other cell clusters combined (T-cells, B-cells, endothelial, myeloid and fibroblast cells). In boxplot, horizontal lines describe median, interquartile range and whiskers extend to 1.5*inter-quartile range. **D)** Barplot displaying the number of TFs (y-axis) passing a Bonferroni adjusted < 0.05 threshold and exhibiting decreased (DN) or increased activity (UP) in tumor epithelial cells (SCIRA & VIPER-D) indicated in darkgreen and darkred, respectively, and correspondingly the same numbers for differential expression (DE). P-values are from a Binomial-test, to test if there is a skew towards inactivation/downregulation in cancer. Right panel depicts the Monte-Carlo (n=1000 runs) significance analysis with grey curve denoting the null distribution for the fraction of TFs exhibiting significant inactivation in tumor epithelial cells, and darkgreen line labeling the observed fraction (0.92=35/38). Empirical P-value derived from the 1000 Monte-Carlo runs is given. **E)** Scatterplot as in A), but now with cells color-labeled according to activation of *FOXA2* as estimated using SCIRA. Beanplots of the predicted SCIRA activity level of *FOXA2* between normal alveolar, tumor epithelial and all other cells. P-value is from a t-test between normal alveolar and tumor epithelial cell clusters. **F)** Pattern of differential activity (SCIRA & VIPER-D) and differential expression for the 38 lung-specific TFs. Darkgreen denotes significant inactivation or underexpression in tumor epithelial cells compared to normal alveolar, brown denotes significant activation or expression. Grey=no-change (NC) and white indicates missing regulon information (VIPER-D). **G)** Pattern of differential expression for the same 38 lung-specific TFs in the bulk RNA-Seq lung cancer datasets (LUSC=lung squamous cell carcinoma, LUAD=lung adenoma carcinoma). **H)** Barplot displaying the number of lung-specific TFs displaying significant and directionally consistent changes in both single-cell and bulk RNA-Seq datasets. In the single-cell data we use differential activity for SCIRA and VIPER-D, whereas for DE we use differential expression.

To test the generality of our observations, we next considered a scRNA-Seq study profiling normal colon epithelial cells and tumor colon epithelial cells ^55^. We first used SCIRA to derive a colon-specific regulatory network from GTEX, resulting in 56 colon-specific TFs and associated regulons (**SI table S6, Supplementary File 1**). This list included many well known intestinal factors such as the enterocyte differentiation factors *CDX1*/*CDX2* ^56^, the crypt epithelial factor *KLF5* ^57^ and the intestinal master regulator *ATOH1* ^58,59^. Next, we obtained TF-activity (TFA) estimates for all 56 colon-TFs across a total of 432 single cells (160 normal epithelial + 272 cancer epithelial, C1 Fluidigm) from 11 different colon-cancer patients. Hierarchical clustering over this TFA-matrix revealed clear segregation of single cells by normal/cancer status and not by patient (**Fig.4A**). Of the 56 TFs, 23 exhibited differential activity (Bonferroni P<0.05) with the great majority (87%, 20/23) exhibiting inactivation, indicating a strong statistical tendency for inactivation in cancer cells (Binomial test, P-3e-5, **Fig.4B**). Once again, had we relied on TF-expression itself, no segregation of single-cells by normal/cancer status was evident (**Fig.4A**), and only 13 TFs were differentially expressed (Bonferroni P<0.05) with no obvious trend towards underexpression in cancer (Binomial test, P=0.13, **Fig.4B**). Of note, while *CDX1* and *CDX2* were found to be both inactivated and underexpressed, several TFs like *KLF5* or *ATOH1* with established tumor suppressor roles in colorectal cancer ^60,61^, were only found inactivated via SCIRA (**Fig.4C**). Interestingly, using VIPER-D there was only moderate correlation with SCIRA’s predictions, with VIPER-D not predicting preferential inactivation and failing to predict inactivity of established tumor suppressors like *KLF5* and *CDX1* (**Fig.4B**). Performing the analysis on a per-patient level and focusing on the 3 patients with the largest numbers of both normal and tumor epithelial cells, revealed a similar skew towards inactivation with 8, 15 and 21 TFs exhibiting significantly lower activity across cancer cells (**Fig.4D**), and with effectively no TF exhibiting increased activity. For several TFs and for each of the 3 patients, inactivation events were seen across most if not all cancer cells (**Fig.4D**): for instance, this was the case for *ATOH1,* or the autophagy inducer *TRIM31* ^62^, thus implicating disruption of this novel and specific autophagy pathway in colon cancer ^63^. Using the 5 patients with both normal and cancer cells profiled, we estimated the frequency of inactivation of all 56 colon-specific TFs across the 5 patients, which revealed that *CDX2* and *TRIM31* were inactivated in 80% of the patients, whilst *ATOH1, HNF4A, CDX1* and *TBX10* were inactivated in 60% (**SI fig.S20**).

**Figure-4:**
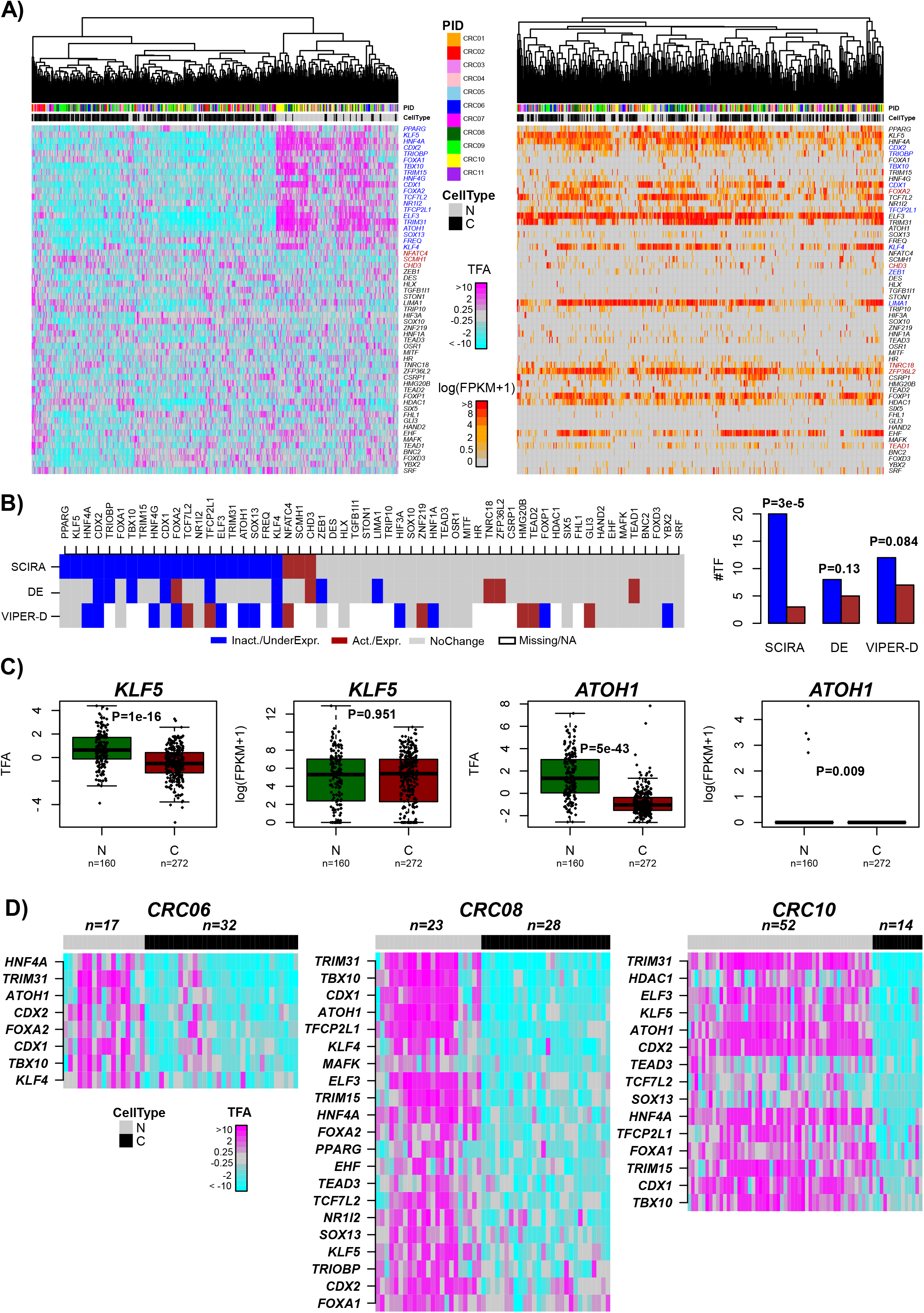
Inactivation of colon-specific TFs in colorectal cancers at single-cell resolution. **A)** Heatmaps of TF-activity (left panel) and TF expression (right panel), with cells ordered by hierarchical clustering over the 56 colon-specific TFs. TFs undergoing significant inactivation/underexpression in cancer cells are labeled in blue, whilst those undergoing activation/overexpression are labeled in darkred. **B)** Heatmap of differential TF-activity (SCIRA & VIPER-D) and TF-expression (DE) between cancer and normal cells, with colors indicating statistical significance (Bonferroni P<0.05) and directionality of change: blue=significant inactivation/underexpression in cancer, brown=significant activation/overexpression in cancer, grey=no-change. Barplots compare the number of inactivated/underexpressed (blue) TFs to the number that are activated/overexpressed (brown). P-values derive from a one-tailed Binomial test to assess significance of skew. **C)** Boxplots displaying TF-activity and TF-expression between normal epithelial and cancer cells for two representative TFs where there is substantial discordance between differential activity and differential expression. P-values for differential TF-activity and TF-expression derive from a t-test and a Wilcoxon rank sum test, respectively. **D)** Heatmaps of TF-activity for the normal and cancer cells from each of 3 patients, and displaying only the subset of the 56 colon-TFs which exhibit significant inactivity in the cancer cells (Bonferroni P < 0.05).

### Tissue-specificity of TF inactivation in cancer

The observed frequent inactivation of tissue-specific TFs in corresponding single cancer cells suggests that these could be driver events. To obtain supporting evidence for this, we reasoned that TFs specific for other unrelated tissue-types would exhibit much lower frequencies of inactivation. We thus compared the lung and colon-specific TFs to additional TFs specific to skin and brain, two non-epithelial tissue types, as well as to stomach-specific TFs which should bear more resemblance to colon-TFs. Consistent with our expectation, in the case of lung cancer cells, the TFs specific to colon, stomach, brain and skin exhibited much lower frequencies of inactivation compared to lung-TFs (**SI fig.21A**). In the case of colon cancer cells, colon and stomach-specific TFs exhibited the highest inactivation frequencies, and were about two-fold higher than for skin and brain-specific TFs (**SI fig.S21B**).

### Inactivation of tumor suppressors in normal cells at risk of cancer

An important application of SCIRA is to normal cells at risk of cancer, which could reveal early inactivation of key tumor suppressor TFs. To demonstrate this, we applied SCIRA to a scRNA-Seq dataset (CEL-Seq) encompassing 564 lung epithelial cells, obtained from bronchial brushings of 6 healthy individuals (6 never-smokers, 6 current smokers) ^64^ (**Methods**). We inferred regulatory activity profiles for our 38 lung-specific TFs in each of the 564 lung epithelial cells, and subsequently used t-stochastic neighborhood embedding (tSNE) ^65^ for dimensional reduction and visualization, as well as DBSCAN ^66^ for clustering (**Methods**), which revealed two main clusters (**Fig.5A**). Overlaying the transcription factor activity (TFA) profiles over the cells revealed that *FOXJ1* (a marker for ciliated cells) was significantly more active in the smaller cluster, suggesting that this cluster defines ciliated cells (**Fig.5A**). Confirming this, *FOXJ1* expression was also higher in this cluster, whilst expression of basal (*KRT5*), club (*SCGB1A1*) and goblet (*MUC5AC*) markers were higher in the larger cluster, suggesting that this larger cluster is composed of non-ciliated lung epithelial cells (i.e. basal cells, goblets and club cells) (**Fig.5B**). Of note, *FOXJ1* was one of the few transcription factors for which activity and expression were reasonably well correlated. For instance, *TBX4* exhibited higher regulatory activity in non-ciliated cells (**Fig.5A**), yet it exhibited a 100% dropout rate across all lung epithelial cells (**Fig.5C**). Other key lung-specific TFs with very high expression in lung tissue, as assessed in our GTEX bulk RNA-Seq data, but with 100% dropout rates included *GATA2* and *TBX2* (**Fig.5C**). Thus, SCIRA is able to retrieve biologically relevant variation in regulatory activity of key TFs, when expression alone can not.

**Figure-5:**
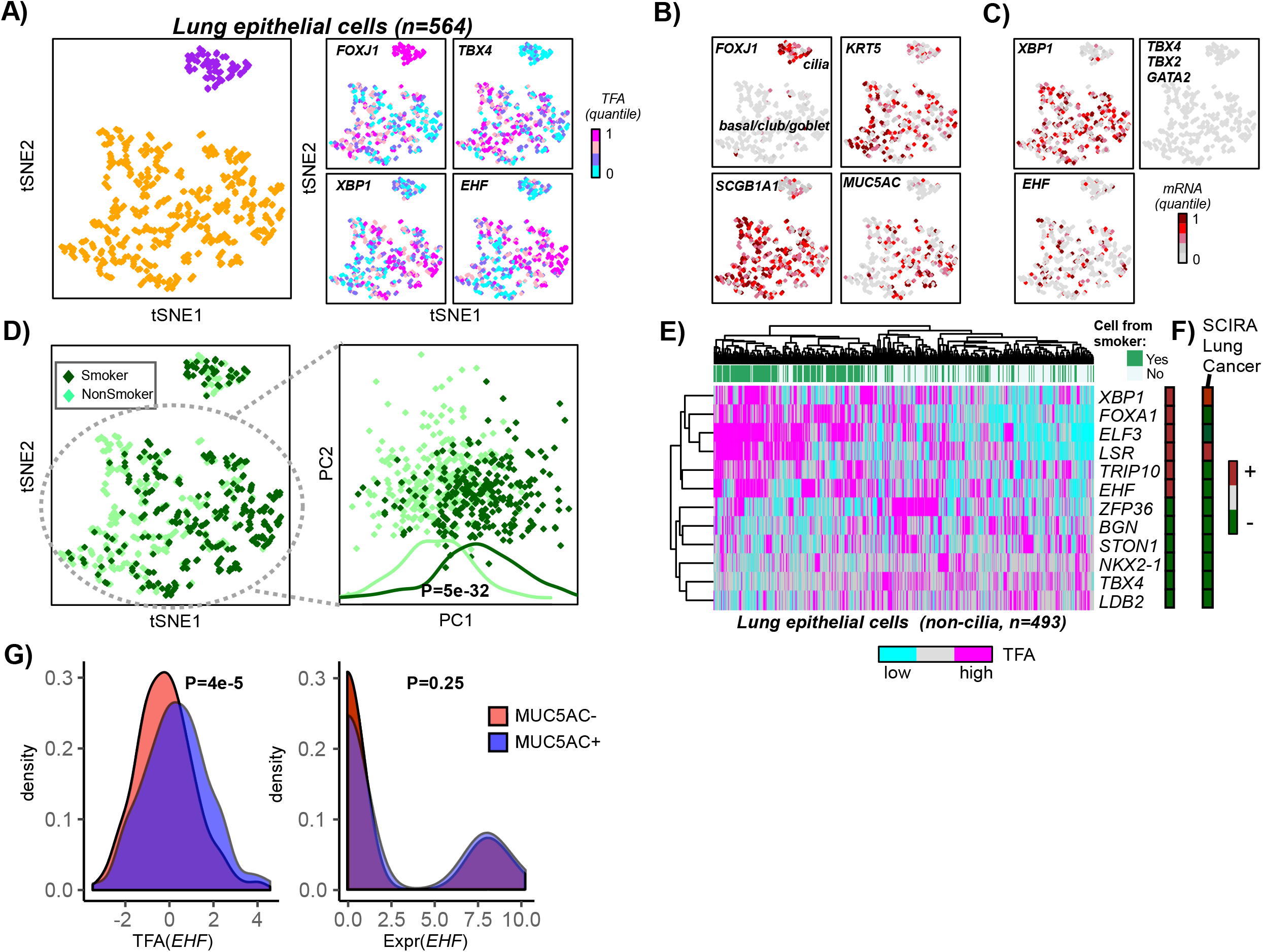
SCIRA reveals smoking-associated tumor suppressor events. **A)** tSNE diagrams of normal lung epithelial cells obtained by application to the SCIRA-derived regulatory activity estimates for the 38 lung-specific TFs. Left panel depicts the two main clusters inferred using DBSCAN, whilst right panels depict the TF-activity levels for 4 of the lung-specific TFs. **B)** As A), but now displaying the mRNA expression levels of 4 markers, one for each of ciliated cells (*FOXJ1*), goblet cells (*MUC5AC*), club cells (*SCBG1A1)* and basal cells (*KRT5*). **C)** As B), but now for 5 lung-specific TFs. **D)** Left panel: As A), but now with cells color-labeled according to whether they derived from a smoker or non-smoker. Right panel: PCA scatterplot (PC1 vs PC2) obtained from a PCA on all non-ciliated cells, plus associated density plots along PC1 for cells stratified according to smoking status. P-value is from a two-tailed Wilcoxon rank sum test. **E)** Hierarchical clustering heatmap over 12 lung-specific TFs exhibiting significant (Bonferroni adjusted P<0.05) activity changes according to smoking-status. Color bar to the right indicates whether TF is more or less active in cells exposed to smoking. **F)** Color bar indicating the pattern of differential regulatory activity for the same 12 TFs in lung cancer cells. **G)** Density distribution of *EHF* activity (left) and *EHF* expression (right) for cells expressing *MUC5AC* (*MUC5AC*+), a goblet cell marker, and cells not expressing *MUC5AC* (*MUC5AC*-). P-values derive from a two tailed Wilcoxon rank sum test.

Despite the tSNE diagram being derived from the regulatory activity profiles of only 38 lung-specific TFs, the larger cluster of non-ciliated cells revealed clear segregation of cells according to whether they derived from current or never-smokers, suggesting that smoking exposure has a dramatic effect on the regulatory activity of lung-specific TFs (**Fig.5D**). We verified this by applying PCA to the activity profiles over the non-ciliated cells only (Wilcox test P=5e-32, **Fig.5D**). We identified a total of 6 TFs exhibiting significantly lower and 6 exhibiting significantly higher regulatory activity in the cells of smokers (**Fig.5E**). Interestingly, among the 6 TFs exhibiting lower activation in cells from smokers, all 6 were also seen to be inactivated in single lung cancer cells, whilst 2 of the 6 exhibiting activation in exposed cells also exhibited increased activity in lung cancer (**Fig.5F**). Among the 6 TFs exhibiting lower activity in both lung epithelial cells of smokers and cancer patients, it is worth noting *NKX2-1*, a putative tumor suppressor for lung cancer as noted recently ^48^, and *TBX4*, another putative tumor suppressor for non-small cell lung cancer ^67,68^. Among the TFs exhibiting increased regulatory activity in smokers we observed *EHF* (**Figs.5E**), a transcription factor which has been implicated in goblet cell hyperplasia ^69^. Consistent with this, goblet hyperplasia is observed in lung tissue from smokers ^64^, and according to SCIRA *EHF* regulatory activity was correlated with expression of the goblet cell marker *MUC5AC* (**Fig.5G**), whereas *EHF* expression itself was not, highlighting once again that SCIRA can recapitulate biological differential activity patterns not obtainable via TF-expression alone. Given that there is goblet cell expansion in smokers ^64^, the increased regulatory activity of *EHF* and other TFs like *ELF3* in smokers could reflect this increase. Of note *ELF3* becomes inactivated in lung cancer cells (**Fig.5F**), which is consistent with its role in lung epithelial cell differentiation being impaired in cancer ^70^ ^71^.

### SCIRA is scalable to millions of cells

Finally, we note that SCIRA can estimate regulatory activity in a manner that scales linearly with the number of profiled cells, thus making it easily scalable to scRNA-Seq studies profiling 100s of thousands to a million cells. In the application to the kidney DropSeq dataset (**SI table S5**) which profiled 9190 cells, runtime was under 4 minutes for 4 processing cores, and under 10 minutes with the regulon-inference step in GTEX included. We performed a subsampling analysis on the kidney set, recording runtimes for manageable numbers of cells, fitted linear functions on a log-log scale, and subsequently estimated runtimes for larger scRNA-Seq studies profiling up to a million cells (**Methods**). In a scRNA-Seq study profiling one million cells, SCIRA would take approximately 100 minutes on 4-cores, or only 4 minutes on a 100-node HPC, whereas other methods would run for months on the same 100-node HPC (**Fig.2D**). Only VIPER-D exhibited a marginally improved computational efficiency compared to SCIRA (**Fig.2D**), owing to the fact that the TF-regulons are derived from a database and are thus precomputed. Thus, SCIRA offers scalability where most competing methods do not.

## Discussion

Dissecting the cellular heterogeneity of cancer, preinvasive lesions and normal tissue at cancer risk is a critically important task for personalized medicine, and it is clear that mapping such cellular heterogeneity needs to be done at single-cell resolution. In the context of cancer risk prediction, the ability to measure gene expression in single normal cells from individuals exposed to an environmental risk factor, could help identify those at most risk of cancer development. Our rationale was to focus on transcription factors that are important for the specification of a given tissue-type, since there is substantial evidence that inactivation/silencing of these transcription factors is an early event in oncogenesis, present in normal cells at risk of neoplastic transformation and thus preceding cancer development itself ^2–4,7–9,72–74^. It follows that identifying such early “tumor suppressor” inactivation events in normal cells at cancer risk in single-cell data could allow prospective identification of individuals at higher risk of cancer development. As demonstrated here, using scRNA-Seq profiles to identify silencing of tissue-specific TFs lacks sensitivity due to the high dropout rate. Instead, we have presented an alternative strategy called SCIRA, which we have very comprehensively validated on many scRNA-Seq datasets profiling normal cells, demonstrating that it can substantially improve the sensitivity and precision to detect correct dynamic TF-activity changes at single-cell resolution.

Application of SCIRA to two scRNA-Seq datasets profiling both normal and cancer cells revealed preferential inactivation of tissue-specific TFs in the corresponding cancer cells, an important biological and clinical insight, which we would not have obtained had we used differential expression. These results are not only in line with analogous findings obtained in bulk RNA-Seq cancer studies ^5^, but helps to further establish which key tissue-specific TFs are inactivated in cancer epithelial cells independently of changes in stromal composition, which could otherwise confound results. For instance, in a tissue like lung, at least 40% of cells are stromal cells ^75^, and so differential expression changes seen in bulk cancer tissue may not be observed or may not be due to expression changes in the epithelial compartment. On the other hand, some consistency with observations in bulk data should be expected, and in this regard we stress that, unlike SCIRA, differential expression approaches on single-cell data did not reveal any consistent patterns with those observed at the bulk level. This inconsistency between single cell and bulk differential expression in cancer is therefore another important insight which demonstrates the need and added value of SCIRA to uncover key tumor suppressor events. For instance, many of the lung-specific TFs which SCIRA predicts to be inactivated in lung tumor epithelial cells (e.g. *NKX2-1, FOXA2, FOXJ1*, *AHR, HIF3A*) ^30^ implicate key cancer-pathways (lung development, alveolarization, ciliogenesis, immune-response, hypoxia-response), and their inactivation likely represent key driver events. Supporting this, epigenetically induced silencing of *NKX2-1* has been proposed to be a key driver event in the development of lung cancer ^48,76^. In the case of colon, our results in the scRNA-Seq data confirm a tumor suppressor role for TFs like *CDX1/CDX2* ^77^, but also serve to reinforce a novel putative tumor suppressor role for *ATOH1* ^78^, for the autophagy inducer *TRIM31* ^62^ and *KLF5* ^79^. Of note, these last three TFs did not exhibit clear significant differential expression changes, yet they were highly significant via analysis with SCIRA.

In the application to normal lung cells from smokers and non-smokers, no preferential inactivation of lung-specific TFs in smokers was observed, consistent with observations derived from buccal (squamous epithelial) cells ^8^. This would suggest that in normal cells exposed to a risk factor, such inactivation events may not yet be under significant selection pressure, yet some of the inactivation events, if present, could be important indicators of future cancer risk. In line with this, out of the 6 lung-specific TFs that were observed to be inactivated in normal lung cells from smokers, all 6 were also inactivated in lung cancer cells. This list included *NKX2-1* and *TBX4*, both of which have tumor suppressor functions ^67,76^. We also observed 6 lung-specific TFs exhibiting increased regulatory activity in cells from smokers, which included *ELF3, XBP1* and *EHF.* Interestingly, *EHF* has been implicated as a driver of goblet hyperplasia ^69^, which is observed in the lung tissue of smokers ^64^. Our data supports the view that *EHF* is a marker of goblet cells and that the increased expression in smokers could be due to an increase in relative goblet cell numbers as observed by Duclos ^64^. Whilst *ELF3* has been reported to be a tumor suppressor in many epithelial cancer types, its function has also been observed to be highly cell-type specific with reported oncogenic roles in lung adenoma carcinoma (LUAD) ^80^. Here we observed *ELF3* activation in the lung non-ciliated cells from smokers and overexpression in bulk lung squamous cell carcinoma (LUSC) tissue, but inactivation in single lung cancer cells (predominantly LUAD) and no expression change in bulk-tissue LUAD. Thus, in future it will be important to profile larger numbers of cells in the lung epithelial compartment of healthy smokers and non-smokers, including lung cancer patients from LUAD, LSCC, NSCLC subtypes, to determine if differential activity patterns are specific to individual lung-epithelial cell subtypes.

Here, and due to obvious limitations on data availability at single-cell resolution, we could not assess the specific mechanism associated with tissue-specific TF silencing in cancer. However, in the context of bulk-tissue data from the TCGA, we have previously shown that the preferential silencing of tissue-specific TFs in cancer is predominantly associated with promoter DNA hypermethylation ^5^. Indeed, inactivation through somatic mutation or copy-number loss/deletion is not a frequent event when considering tissue-specific TFs ^5^, in contrast to other gene-families like kinases, epigenetic enzymes or membrane receptors which do exhibit more frequent genetic alterations ^81,82^. Thus, it is very likely that the observed inactivation of tissue-specific TFs in individual cancer cells is also associated with promoter DNA hypermethylation.

In summary, we have presented and validated a computational strategy called SCIRA that can improve the sensitivity and precision to detect regulatory activity changes of key tissue-specific transcription factors in scRNA-Seq data, and that can reveal tumor suppressor events at single-cell resolution which would otherwise not be possible using differential expression. SCIRA has shown that tissue-specific TFs are preferentially inactivated in corresponding cancer cells, suggesting that these could be tumor suppressor driver events. Importantly, SCIRA also provides a scalable framework in which to infer tissue-specific regulatory activity in scRNA-Seq studies profiling even millions of cells. We envisage that SCIRA will be particularly useful for scRNA-Seq studies aiming to identify altered differentiation programs in normal tissue exposed to cancer risk factors, preinvasive lesions and cancer at single-cell resolution. This is important as this may offer clues and insight into the earliest stages of oncogenesis.

## Methods

### Single cell data and preprocessing

We analyzed scRNA-Seq data from a total of 6 studies:

#### Lung Differentiation set

This scRNA-Seq Fluidigm C1 dataset derives from Treutlein et al ^35^. Normalized (FPKM) data were downloaded from GEO under accession number GSE52583 (file: GSE52583.Rda). Data was further transformed using a log2 transformation adding a pseudocount of 1, so that 0 FPKM values get mapped to 0 in the transformed basis. After quality control, there are a total of 201 single cells assayed at 4 different stages in the developing mouse lung epithelium, including embryonic day E14.5 (n = 45), E16.5 (n = 27), E18.5 (n = 83) and adulthood (n = 46).

#### Liver Differentiation set

This scRNA-Seq Fluidigm C1 dataset was derived from Yang et al 36, a study of differentiation of mouse hepatoblasts into hepatocytes and cholangiocytes. Normalized (TPM) data was downloaded from GEO under accession number GSE90047 (file: GSE90047-Singlecell-RNA-seq-TPM.txt). Data was further transformed using a log2 transformation adding a pseudocount of 1, so that 0 TPM values get mapped to 0 in the transformed basis. After quality control, 447 single-cells remained, with 54 single cells at embryonic day 10.5 (E10.5), 70 at E11.5, 41 at E12.5, 65 at E13.5, 70 at 14.5, 77 at 15.5 and 70 at E17.5.

#### Pancreas Differentiation set

This scRNA-Seq Smart-Seq2 data derives from Yu et al ^37^, profiling single cells during murine pancreas development, from embryonic stages E9.5 to E17.5. Normalized (TPM) data was downloaded from GEO (GSE115931, file: GSE115931_SmartSeq2.TPM.txt”). Data was further log2-transformed with a pseudocount of 1. After quality control, 2195 cells remained: 113 (E9.5), 211 (E10.5), 263 (E11.5), 252 (E12.5), 421 (E13), 338 (E14.5), 242 (E15), 185 (E16.5), 170 (E17.5).

#### Kidney-organoid Differentiation set

This scRNA-Seq DropSeq data derives from Wu et al ^38^, profiling single cells in a kidney organoid differentiation experiment (Takasato protocol) starting out from iPSCs, with 218 cells profiled at day-0, 1741 at day-7, 1169 at day-12, 1097 at day-19 and 4965 at day-26. Read count data for all 9190 high quality cells was downloaded from GEO (GSE118184, file: GSE118184_Takasato.iPS.timecourse.txt”). Counts were scaled for each cell by the total read count, multiplied by a common scaling factor of 10^4^ and subsequently log2-transformed with a pseudocount of 1.

#### Normal and cancer lung tissue dataset

This scRNA-Seq 10X Chromium dataset was derived from ^47^, a study profiling malignant and non-malignant lung samples from five patients. We downloaded all .Rds files available from ArrayExpress (E-MTAB-6149), which included the processed data and t-SNE coordinates, as well as cluster cell-type assignments. After quality control, a total of 52,698 single-cells remained of which 1709 were annotated as alveolar, 5603 as B-cells, 1592 as endothelial cells, 1465 as fibroblasts, 9756 as myeloid cells, 24911 as T-cells and 7450 as tumor epithelial cells. A small cluster of 212 cells was annotated as normal epithelial, yet they derived from a malignant sample ^47^, so given this inconsistency we removed these cells from any analysis, as according to us their “normal” nature is far from clear. The alveolar epithelial cell cluster derived mainly from non-malignant samples and was therefore considered most representative of the normal epithelial cells found in lung.

#### Normal and cancer colon dataset

This scRNA-Seq Fluidigm C1 dataset is derived from ^55^, a study profiling malignant and non-malignant colon epithelial cells from 11 patients. We downloaded the normal mucosa and tumor epithelial cell FPKM files from GEO under accession number GSE81861. In total there were 160 and 272 normal and tumor epithelial cells, respectively, as determined by the original publication.

#### Normal lung from smokers and non-smokers

This scRNA-Seq dataset is derived from ^64^, where FACS sorted lung epithelial cells from 6 never-smokers and 6 smokers were analysed with the CEL-Seq platform. We downloaded the raw UMI counts from GEO under accession number GSE131391. We followed a similar normalization and QC procedure as described in 64, although we used a more stringent cell quality criterion, removing any cells with a total UMI count less than 2400. This threshold was chosen because the total UMI count per cell exhibit a natural bimodal distribution, with the value 2400 defining the natural decision boundary between low and high quality cells. This resulted in 564 epithelial cells. For these cells data was further normalized by scaling UMI counts to TPM, adding a pseudocount of 1 and finally taking the log_2_ transformation. We note that results reported here were unchanged if not scaling UMI counts, i.e. if using log_2_(UMI+1).

### Bulk tissue mRNA expression datasets

For applying SCIRA to data from epithelial tissues, we used the bulk RNA-Seq dataset from the GTEX resource ^24^ to infer regulons. Specifically, the normalized RPKM data was downloaded from the GTEX website and annotated to Entrez gene IDs. Data was then log_2_ transformed with a pseudocount of +1. This resulted in a data matrix of 23929 genes and 8555 samples, encompassing 30 tissue types (adipose=577, adrenal gland=145, bladder=11, blood=511, blood vessel=689, brain=1259, breast=214, cervix uteri=11, colon=345, esophagus=686, fallopian tube=6, heart=412, kidney=32, liver=119, lung=320, muscle=430, nerve=304, ovary=97, pancreas=171, pituitary=103, prostate=106, salivary gland=57, skin=891, small intestine=88, spleen=104, stomach=193, testis=172, thyroid=323, uterus=83, vagina=96). In addition, we also analyzed the bulk RNA-Seq dataset from the lung TCGA studies ^53,54^, which was normalized as described in our previous publications ^5,83^.

### The SCIRA algorithm

The SCIRA algorithm has two main steps: (i) construction of a tissue-specific regulatory network and (ii) inference of regulatory activity in single cells for the transcription factors (TFs) in the network constructed in step (i).

#### (i) Construction of tissue-specific regulatory network

For a given tissue-type, SCIRA infers a corresponding tissue-specific regulatory network using a greedy partial correlation algorithm framework called SEPIRA ^8^. The greedy partial correlation approach is similar in concept to the GENIE3 algorithm ^84^ (which was found to be one of the best performing reverse-engineering methods in the DREAM-5 challenge ^85^), in the sense that it infers the candidate regulators for each gene in turn. However, we use partial correlations instead of regression trees. By computing partial correlations over the GTEX dataset, which consists of 8555 samples across 30 different tissue-types, it is possible to identify direct regulatory relations that are relevant in the context of differentiation and development. Briefly, having log-transformed the GTEX RNA-Seq set, as described previously ^8^, we first select genes with a standard deviation larger than 0.25, so as to remove genes with no significant expression variation across the 8555 samples. A total of 19478 genes with Entrez gene annotation were left after this step. Next, we used a list of 1385 human TFs as defined by the TRANSC_FACT term of the Molecular Signatures Database ^86^, of which 1313 had representation in our filtered GTEX set. Genes not annotated as TFs, were considered putative targets, and we first estimated Pearson correlations between the 1313 TFs and the 18165 targets. Using a conservative P-value threshold of 1e-6 to define putative interactions between TFs and targets, we next selected TFs with at least 10 putative targets. For each target-gene *g* and its putative TF regulators *f,* we then computed partial correlations between *g* and *f,* as

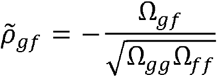

where Ω is the inverse of the expression covariance matrix, which is of dimension (1+*nf*)*(1+*nf*) with *nf* the number of putative TF regulators. Importantly, by estimating the partial correlations in a greedy fashion, i.e. for each target gene separately, the inverse of the covariance matrix is always well defined (no need to estimate a pseudo-inverse) since *nf* ≪ 8555, i.e much less than the number of samples over which the partial correlations are estimated. In other words, we estimate the partial correlations between each target gene and its candidate regulators from the marginal analysis above, and we do this for each target gene separately, which thus provides a natural regularization. Partial correlation thresholds of +/− 0.2, or even +/− 0.1 are statistically significant given the large number of samples (8555) in the GTEX set (as verified by random resampling), so we use either one of these thresholds depending on the number of TFs desired, although the number of resulting TFs is similar for both choices of threshold. This then defines a global regulatory network between TFs and target genes, where indirect dependencies have been removed due to the use of partial correlations ^87^.

The final step is the construction of a tissue-specific regulatory network as the subnetwork obtained by identification of tissue-specific TFs, i.e. TFs with significantly higher expression in the given tissue type compared to all other tissues combined. This is done using the empirical Bayes moderated t-test framework (limma) ^88^. Importantly, a second limma analysis is performed by comparing the tissue of interest to individual tissue types if these contain cells that are believed to significantly infiltrate and contaminate the tissue of interest.

Thus, in the case of liver we perform two limma analyses: comparing liver to all other tissue-types, and separately, liver to only blood and spleen combined, since blood/spleen consists of immune cells which are known to infiltrate liver tissue accounting for approximately 40% of all cells found in liver ^75^. We require a liver-specific TF to be one with significantly higher expression in both comparisons: when comparing to all tissues we use an adjusted P-value threshold of 0.05 and a log2(FC) threshold of log2(1.5)≈0.58, whereas when comparing to blood/spleen we only use an adjusted P-value threshold of 0.05. This ensures that the identified TFs are not driven by a higher immune cell (IC) infiltration in the tissue of interest compared to an “average” tissue where the IC infiltration may be low. As applied to liver and using a significance threshold on partial correlations of +/− 0.2, SCIRA/SEPIRA inferred a network of 22 liver-specific TFs and their regulons, with the average number of genes per regulon being 41, and with range 10 to 151. This network is available as an Rds file “netLIV.Rds” in **Supplementary File 1**. The same procedure was used for the other tissue-types and the corresponding networks for pancreas (netPANC.Rds), kidney (netKID.Rds) and colon (netCOL.Rds) are also available in **Supplementary File 1.**

We note that regulon genes could be selected further based on whether they are direct binding targets of the TF, as for instance determined by a ChIP-Seq assay. However, we did not pursue this strategy here, for a number of good reasons. First, the definition of a regulon, as originally proposed by Andrea Califano’s lab ^31,89^, does not require a member of the regulon to be a direct target of the regulator. Indeed, it could well be that a downstream gene in the pathway is an equally good if not even better marker of upstream regulatory activity. Thus, it makes sense to keep all inferred regulon genes in the regulon, following previous studies. On the other hand, some enrichment for direct targets is to be expected, and we indeed checked enrichment for ChIP-Seq binding targets using data from the ChIP-Seq Atlas ^32^. A second reason is that reducing the number of regulon genes also means a loss of power, specially so if the regulon genes are bona-fide markers of upstream regulatory activity. Third, ChIP-Seq data is still very limited in the number of cell-types profiled, which may not include a representative cell-type of the tissue in question. In other words, the sensitivity of a ChIP-Seq assay is also limited and if a gene is not predicted to be a binding target in cell-type “A” it could still be a direct target in the tissue/cell-type of interest.

#### (ii) Estimation of regulatory activity

Having inferred the tissue-specific TFs and their regulons, we next estimate regulatory activity of the TFs in each single cell of a scRNA-Seq dataset. This is done by regressing the log-normalized scRNA-Seq expression profile of the cell against the “target-profile” of the given TF, where in the target profile, any regulon member is assigned a +1 for activating interactions, a −1 for inhibitory interactions. All other genes not members of the TF’s regulon are assigned a value of 0. The TF-activity is then defined as the t-statistic of this linear regression. Before applying this procedure the normalized scRNA-Seq dataset is z-score normalized, i.e. each gene is centered and scaled to unit standard deviation.

We note that SCIRA relies on the tissue-specific regulatory network inferred in step-1. As such, SCIRA is particularly useful for scRNA-Seq studies that profile cells in the tissue of interest, either as part of a developmental or differentiation timecourse experiment, or in the context of diseases where altered differentiation is a key disease hallmark e.g. cancer and precursor cancer lesions.

### Pseudocode implementing SCIRA algorithm

The previously described steps implementing SCIRA can be run using the functions provided as part of the SEPIRA Bioconductor package, or preferably from the SCIRA-package: https://github.com/aet21/scira. Briefly, assuming the normalized GTEX RNA-Seq dataset matrix is stored in an R-object called “*data.m*”, with rows labeling genes and columns labeling samples, and assuming we choose liver as our tissue of interest, we would run the following set of commands in order to construct the liver-specific regulatory network:

➢ *net.o* <- *sciraInfReg(data=data.m, sdth=0.25, sigth=1e-6, pcorth=0.2, spTH=0.01, minNtgts=10, ncores=4)*
➢ *livernet.o* <- *sciraSelReg(net.o, tissue=colnames(data.m), toi=”Liver”, cft=”Blood”, degth=c(0.05,0.05), lfcth=c(log2(1.5),0))*

In the above *colnames(data.m)* labels the tissue-type of each sample (column) of the data matrix. Note that the parameter *cft* labels the confounding tissue-type, which in this case is blood, because immune-cells, the main component of blood, is a major contaminant cell-type in liver-tissue ^75^. One important parameter in the above function, which directly controls the number of retrieved TFs is *spTH:* this parameter controls the number of significant correlations in the marginal analysis to be included in the subsequent partial correlation analysis. By default this is set at 1% of all possible interactions, but increasing this threshold to 5 or 10% will increase the number of interactions and thus the number of retrieved TFs. The tissue-specific regulatory network can be found in the *livernet.o$netTOI* entry, which is a matrix with columns labeling the tissue-specific transcription factors and rows labeling gene targets. The entries in this matrix are either 1 for a positive interaction, 0 for no interaction, and −1 for inhibitory associations. This matrix provides the regulons to the function for estimating regulatory activity in a bulk sample or in single-cells. For instance, assuming that we have a log-normalized scRNA-Seq dataset representing liver development in humans, *scRNA.m*, we would obtain regulatory activity estimates for each of the transcription factors present in *livernet.o$netTOI,* by running:

➢ *actTF.m <- sciraEstRegAct(data=scRNA.m, regnet=livernet.o$netTOI, norm=”z”,ncores=4)*

where the *norm* argument specifies that genes in the *scRNA.m* data matrix should be z-score normalized, before estimating regulatory activity. We note that the output object *actTF.m* would define a matrix with rows labeling the tissue-specific transcription factors and columns labeling the single-cells, and with matrix entries representing regulatory activities. We further note that the tissue-specific regulatory networks derived from GTEX, as used in this work, are provided in **Supplementary File 1.** Full details of how to run scira are provided in the vignette of the scira R-package.

### Power-calculation for SCIRA

We derived a formula to estimate the sensitivity (which we shall denote by *SE*) of SCIRA to detect highly expressed cell-type specific TFs in a given tissue, as a function of the corresponding cell-type proportion in the tissue. The main parameters affecting the power estimate include the relative sample sizes of the two groups being compared (*n*_*1*_ and *n*_*2*_), the average expression effect size *e* (in effect the average expression fold-change) of the cell-type specific TFs compared to all other cell-types, which will depend on the proportion of the cell-type (*w*) within the tissue of interest. Indeed, it is not difficult to prove that under reasonable assumptions ^90^, the sensitivity (*SE*) is given by the formula

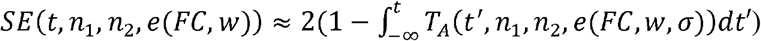

where *t* is the statistic value (we assume a t-statistic) dictating the significance threshold, and where *T*_*A*_ denotes the non-central Student’s t-distribution with non-centrality parameter *μ* equal to

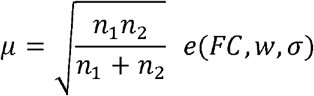

We note that the effect size *e* is of the form 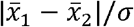, i.e. the ratio of the difference in average expression between the two groups divided by a common pooled standard deviation that reflects the intrinsic variance in each group. We note that we are assuming that the bulk RNA-Seq data has been log-normalized so that *e* is derived from the log-transformed data. For instance, if a gene (say a TF) shows the same expression distribution for all cell-types in the tissue of interest compared to all other tissues, then 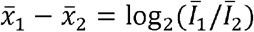 where 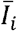 denotes the average intensity (i.e. FPKM/TPM) value in group-*i.* Assuming that the given TF is only more highly expressed in a cell-type that makes up only a proportion *w* of the cells in the tissue of interest, then *e* = log_2_[*FC* * *w* + 1 * (1 − *w*)]/*σ* where *FC* is the average fold-change. To estimate the sample sizes for the power calculation, we note that the median number of samples per tissue-type in GTEX is approximately 170. We took a more conservative value of *n*_*1*_=150 to represent the number of samples in our tissue of interest, with the rest of samples in GTEX, i.e. *n*_*2*_=8555−150=8405, defining the number of samples from other tissue-types. To estimate the average expression fold-change *FC* for top DEGs between single-cell types in a tissue, we analysed expression data from purified FACS sorted luminal and basal cells from the mammary epithelium ^91^. Because FACS sorted cell populations are still heterogeneous, we thus expect the resulting fold change estimates to be conservative. Using limma ^88^, we estimated *FC* to be 8 for the highest ranked DEGs, and approximately 6 for the top 200-300 DEGs. We note that these estimates are for a scaled basis where σ=1. Thus, we approximate the effect size *e* ≈ log_2_[*FC* * *w* + 1 * (1 − *w*)] with *FC=8* or *6,* so as to consider two different effect size scenarios. For the proportion *w* we assumed two values: *w*=*0.05* and *w*=*0.2* representing 5 and 20% of the cells in the tissue of interest. Note that if *w*=*1*, all cells within the issue of interest exhibit differential expression at magnitude *FC* and if *w*=*0*, no cell is differentially expressed. Finally, to compute the sensitivity as a function of the significance level threshold *t*, we used the parameters above as input to the TOC function of the OCplus R-package ^90^.

### Implementation of scImpute, MAGIC and SCRABBLE

scImpute (version 0.0.9) ^39^ was run with default parameters (labeled =FALSE, drop_thre=0.5) in all analysis, with the exception of the Kcluster parameter, which was chosen to reflect the number of underlying cell-types in each tissue analysed: Liver=3, Lung=4, Pancreas=15, Kidney= 14, i.e this parameter was set for each tissue following the numbers of cell-types as specified in the original papers. For MAGIC (version 1.4.0) ^40^ in liver, lung and pancreas, we used the following parameters: k = 15, alpha= 5, t = “auto”, knn_dist.method= “euclidean”. For kidney, because of the much larger number of cells, we chose larger values for k=30 and alpha=10. The number of PCs (npca) was determined in all tissues as the number of PCs explaining 70% of variation in the data, as recommended ^40^. For SCRABBLE (version 0.0.1) ^41^, the average bulk RNA-seq expression vector was computed using the corresponding tissue-type samples from the GTEX dataset. The alpha parameter in the function was chosen for each tissue-type, following the recommendations given in the paper: Liver=1, Lung=1, Pancreas=0.1, Kidney=0.1. The other parameter values were beta = 1e-5, gamma = 0.01. For all other parameters, we used the default choices: nIter = 20, error_out_threshold = 1e-04, nIter_inner = 20, error_inner_threshold = 1e-04.

### Implementation of GENIE3 and SCENIC

SCENIC is a pipeline of 3 distinct methods (GENIE3, RcisTarget, AUCell), each with its own Bioconductor package. We used the following versions: GENIE3_1.4.0, RcisTarget_1.2.0 and AUCell_1.4.1. Because the lung, liver and pancreas scRNA-Seq sets are from mice, we used as regulators a list of 1686 mouse TF from the RIKEN lab (http://genome.gsc.riken.jp/TFdb/) together with the homologs of the human TFs in our lung, liver and pancreas specific networks if these were not in the RIKEN lab list. GENIE3 was run with default parameter choices (treeMethod=”RF”, K=”sqrt”, nTrees=1000) but on a reduced data matrix where genes with a standard deviation less than 0.5 were removed. Regulons of TFs were obtained from GENIE3 using a threshold on the inferred weights (representing the regulatory strength and termed “importance measure” in GENIE3) of 0.01, and only positively correlated targets were selected using a Spearman correlation coefficient threshold > 0. In SCENIC, the targets are then scanned for enriched binding motifs using RcisTarget. We used the 7species.mc9nr feather files for both 500bp upstream of the TSS and also for a 20kb window centered on the TSS. Any enriched motifs in both analyses were combined to arrive at a single list of enriched motifs and associated TFs. We then found the overlap with the annotated TFs from GENIE3, and only those that overlapped were considered valid TF regulons. For these we then estimated a regulatory activity score using an approach similar to the one implemented in AUCell, but one that is threshold independent, and therefore an improvement over the method used in AUCell. Specifically, the activity score was defined as the AUC of a Wilcoxon rank sum test, whereby in each single cell, genes are first ranked in decreasing order of expression, and the AUC-statistic is then derived by comparing the ranks of the regulon (all positively correlated) genes to the ranks of all other genes.

### Implementation of VIPER-D

In order to assess the importance of the tissue-specific regulons used in SCIRA, we compared SCIRA to a method that uses non tissue-specific TF-regulons. We note that there are tools like PAGODA ^92^ that can infer activity scores from gene sets, yet a regulon also entails directionality (i.e. positive or inhibitory interaction) information, which also needs to be assessed. Hence, motivated by the recent work by Holland et al ^46^, we decided to test SCIRA against the combined use of VIPER ^43^ and the dorothea TF-regulon database ^45^. Of note, VIPER infers regulatory activity in any given sample/cell given a TF-regulon, and that the dorothea TF-regulon database is not tissue-specific, although one of the sources in building dorothea is the same GTEX dataset used by SCIRA to build its tissue-specific regulons. We ran viper with the following argument choices: dnull = NULL, pleiotropy = FALSE, nes = TRUE, method = c("none"), bootstraps = 0, minsize = 5, adaptive.size = FALSE, eset.filter = TRUE, mvws = 1, cores = 4. Dorothea also provides likelihood information that a given regulatory interaction in the database is true, and VIPER allows such likelihood information to be used when inferring regulatory activity. We ran VIPER-D in two ways: (i) assigning the same likelihood to all listed regulatory interactions (ie equal weights), and (ii) by using the likelihood information. In Dorothea, the likelihood is encoded as an ordinal categorical variable: A, B, C, D, E, with A indicating highest confidence. In order to run this with VIPER, we transformed these categories into confidence weights using the mapping: A=1, B=0.8, C=0.6, D=0.4, E=0.2. Results in this manuscript are reported for the case of equal weights.

We note that these likelihoods vary mostly between TFs, and not between the targets of a given TF, which is why results are largely unchanged had we used the likelihood information.

### Differential Expression (DE) analysis

In this work we compare SCIRA to ordinary DE analysis, as implemented using a Wilcoxon rank sum test for binary phenotypes, or using non-parametric Spearman rank correlations for ordinal phenotypes (e.g. multiple timepoints or stages). The use of a non-parametric test, which is distribution assumption free, works well for scRNA-Seq with high dropout rates. When comparing statistics of differential activity from SCIRA to those from DE analysis, we transform Wilcoxon rank sum or Spearman test P-values into z-statistics using a quantile normal distribution, taking into account the magnitude of the AUC value from the Wilcoxon test (i.e. AUC values > 0.5 correspond to higher expression in one group compared to other, whereas AUC < 0.5 represents the opposite case), or the sign of the Spearman correlation coefficient in the case of ordinal phenotypes.

### Comparative sensitivity and precision analysis

We compared SCIRA to seven other methods in their sensitivity and precision to identify gold-standard sets of tissue-specific TFs. These gold-standard sets were constructed from GTEX and validated in orthogonal bulk tissue gene expression datasets from NormalAtlas ^33^ and Roth et al ^34^. The number tissue-specific TFs for liver, lung, pancreas and kidney were 22, 38, 30 and 38, respectively. The seven other methods were ordinary differential DE analysis, scImpute+DE, MAGIC+DE, Scrabble+DE, GENIE3, SCENIC and VIPER-D. We note that SCENIC runs GENIE3 as a first step and then selects TF-regulons for which corresponding TF-binding motifs are enriched. So, for the method denoted “GENIE3” we drop the requirement of TF-binding motif enrichment. For SCIRA, GENIE3, SCENIC and VIPER-D we obtain TF-activity estimates, whereas the other methods rely on direct gene expression, measured or imputed. Sensitivity (SE) was estimated as the fraction of gold-standard TFs which exhibited significant increased activation/expression with differentiation timepoint, as determined using a Bonferroni adjusted P<0.05 threshold. Precision equals 1-FDR (false discovery rate), with the FDR defined by the ratio of significantly inactivated TFs to the total number of significantly differentially active TFs, since inactivation of these TFs is inconsistent with known biology and therefore represent false positives. Correspondingly, for methods relying on differential expression, the FDR is defined by the ratio of significantly downregulated TFs to the total number of significantly differentially expressed TFs.

### Comparative runtime and scalability analysis

Objective comparison of runtimes of the different algorithms is hard because each method has different requirements for input, and because runtimes depend critically on the choice of method-specific parameters. Nevertheless, we compared runtimes for 5 important algorithms (SCIRA, MAGIC, Scrabble, GENIE3/SCENIC and VIPER-D), both in terms of their actual implementations on the liver, lung, and pancreas and kidney sets, but also in a scaling analysis with largely default parameters, where we applied all 5 methods to varying subsets of the kidney scRNA-Seq set (total 9190 cells). Briefly, we processed the scRNA-Seq kidney DropSeq data as described earlier, and filtered genes with sufficient variance resulting in 12596 genes. We then constructed subsets with variable cell numbers by randomly subsampling 200, 400, 600, 800, 1000 and 1500 cells, and ran each of these methods on each of these subsampled datasets. In the case of SCIRA, MAGIC, GENIE3/SCENIC and VIPER-D we ran the algorithms with 4 processing cores on a Dell PowerEdge server with Intel Xeon CPU E5-4660 v4 and clock speed of 2.20HHz. Unfortunately, Scrabble does not offer a parallelizable option and is excruciatingly slow for larger e.g. a 10,000 cell dataset. Thus, for each method, we obtained runtimes as a function of cell-number, and fitted a linear regression to the data on a log-log scale. On a log-log scale where both runtime and cell-number are logged, the relation is generally linear. Next, we imputed runtimes for much larger datasets up to a million cells.

## Data Availability

Data analyzed in this manuscript is already publicly available from the following GEO (www.ncbi.nlm.nih.gov/geo/) accession numbers: GSE52583, GSE90047, GSE115931, GSE118184, GSE81861, GSE131391, and from ArrayExpress (www.ebi.ac.uk/arrayexpress) under accession number E-MTAB-6149.

## Code Availability

The scira R-package is freely available from https://github.com/aet21/scira

## Additional Files

Supplementary File 1 contains R (.Rds) object files, containing the inferred regulatory networks for colon (netCOL.Rds), kidney (netKID.Rds), liver (netLIV.Rds), lung (netLUNG.Rds) and pancreas (netPANC.Rds). Supplementary Information File contains all Supplementary Figures and Supplementary Tables.

## Ethics

All data analyzed in this manuscript is freely available in the public domain, and so no Ethics statement is required as all primary data was already presented elsewhere.

## Competing Interests

The authors declare that there are no competing interests.

## Author Contribution

Study was conceived and designed by AET. Statistical analyses were performed by AET and replicated by NW. Software package was prepared by NW. Manuscript was written by AET.

## Acknowledgements

This work was supported by NSFC (National Science Foundation of China) grants, grant numbers 31571359 and 31771464 and by a Royal Society Newton Advanced Fellowship (NAF award number: 164914). We would also like to thank Peter Kharchenko for useful discussions.

## Notes

### Competing Interest Statement

The authors have declared no competing interest.

