## Supplementary Information for "Improved detection of tumor suppressor events in single-cell RNA-Seq data"

#### SUPPLEMENTARY FIGURES

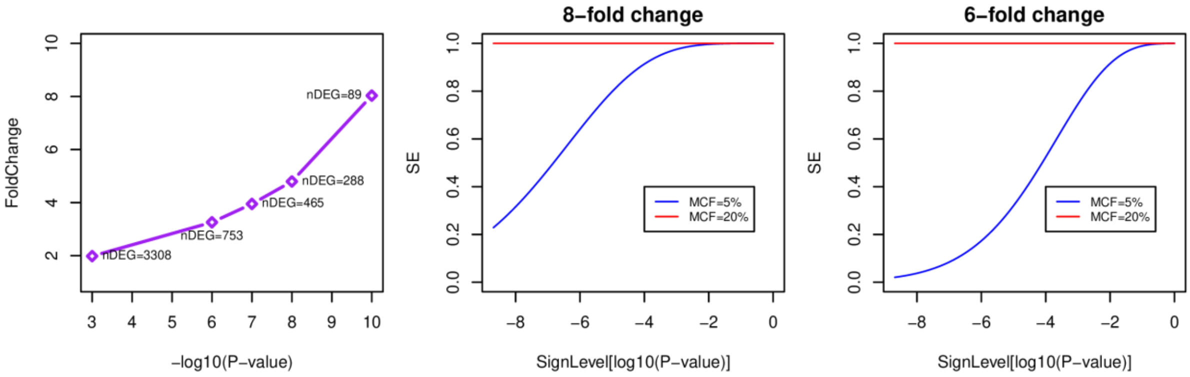


**fig.S1: Fold-change estimation and power-analysis for SCIRA. Left panel:** Estimated Fold-Change (y-axis) between purified FACS sorted luminal and basal bulk samples from the mammary epithelium [1] against the significance-level (-$\log_{10}$[P-value], x-axis) with P-value derived from a moderated t-test. The number of differentially expressed genes (nDEG) at each significance threshold is given. Observe that 6 to 8-fold changes are not uncommon when comparing purified cell populations to each other. **Middle & Right panels**: Sensitivity (SE, y-axis) to detect putative transcription factors exhibiting 8 or 6-fold changes in expression in a cell-type with minor cell fraction (MCF) of 5% and 20% in the tissue (i.e. making up 5 and 20% of the cells in the tissue) against the significance level ($\log_{10}$[P-value], x-axis). The estimation is based using the GTEX dataset where the median number of samples per tissue-type is 150, and where the total number of samples, encompassing 30 tissue-types is 8555. Thus, the power analysis is performed for 150 samples vs 8405. In addition, we have assumed that there are 50 truly differentially expressed TFs within the tissue-type of interest.

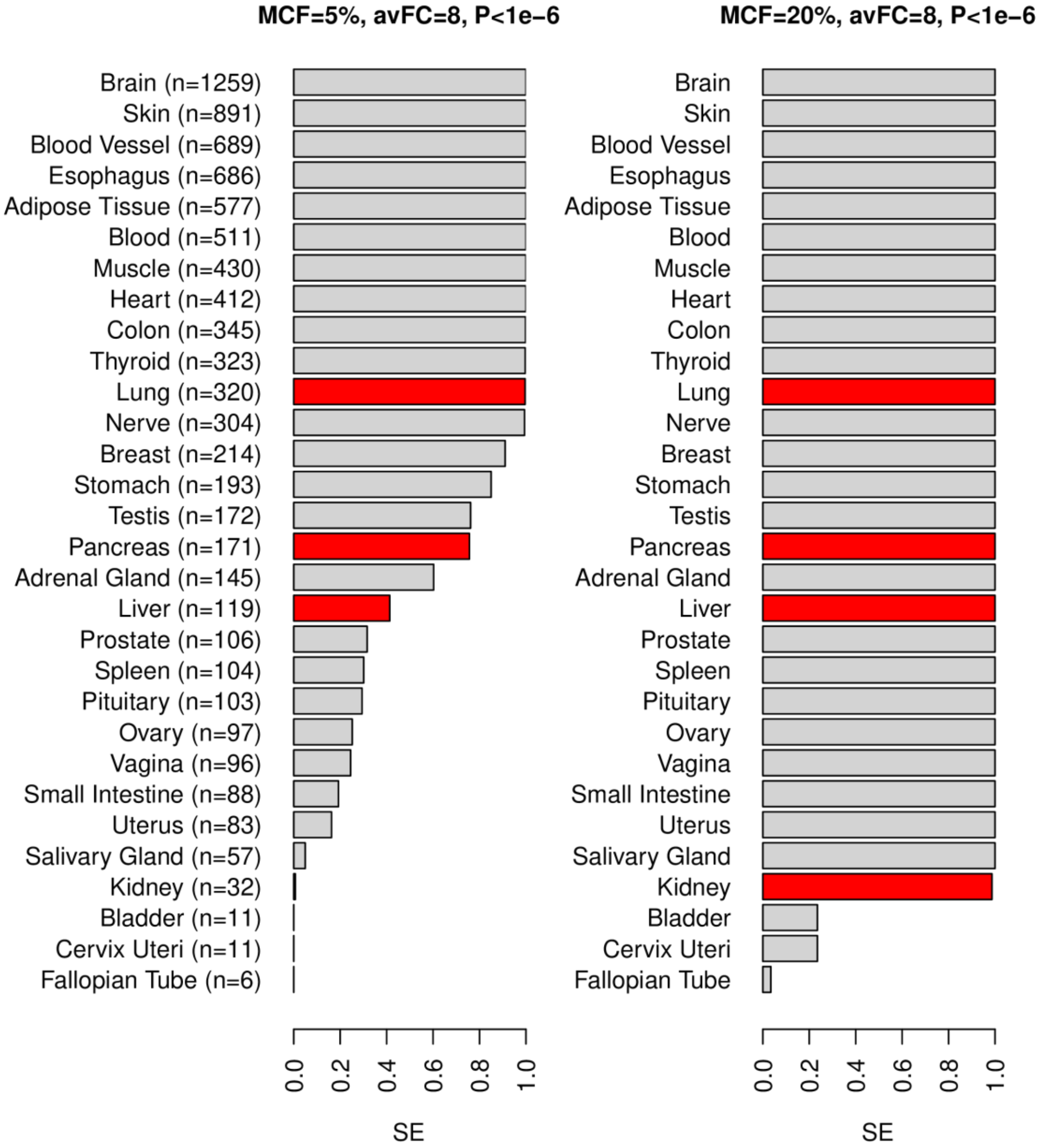


**fig.S2: Power-Analysis for SCIRA in GTEX.** Barplots displaying the sensitivity to detect tissue-specific TFs in each tissue of the GTEX dataset, at a P-value significance threshold of 1e-6, and assuming an average fold-change of 8 (this fold-change estimate is reasonable and has been derived from FACS sorted data), and for two different scenarios: in one case, the TFs are assumed to be overexpressed only in a cell subtype that makes up 5% of the tissue (MCF=5%, left panel), whereas in the other case the cell subtype fraction is assumed to be 20% (right panel). The number of samples in each tissue is indicated in the left panel. In red, we highlight those tissues considered in this manuscript.

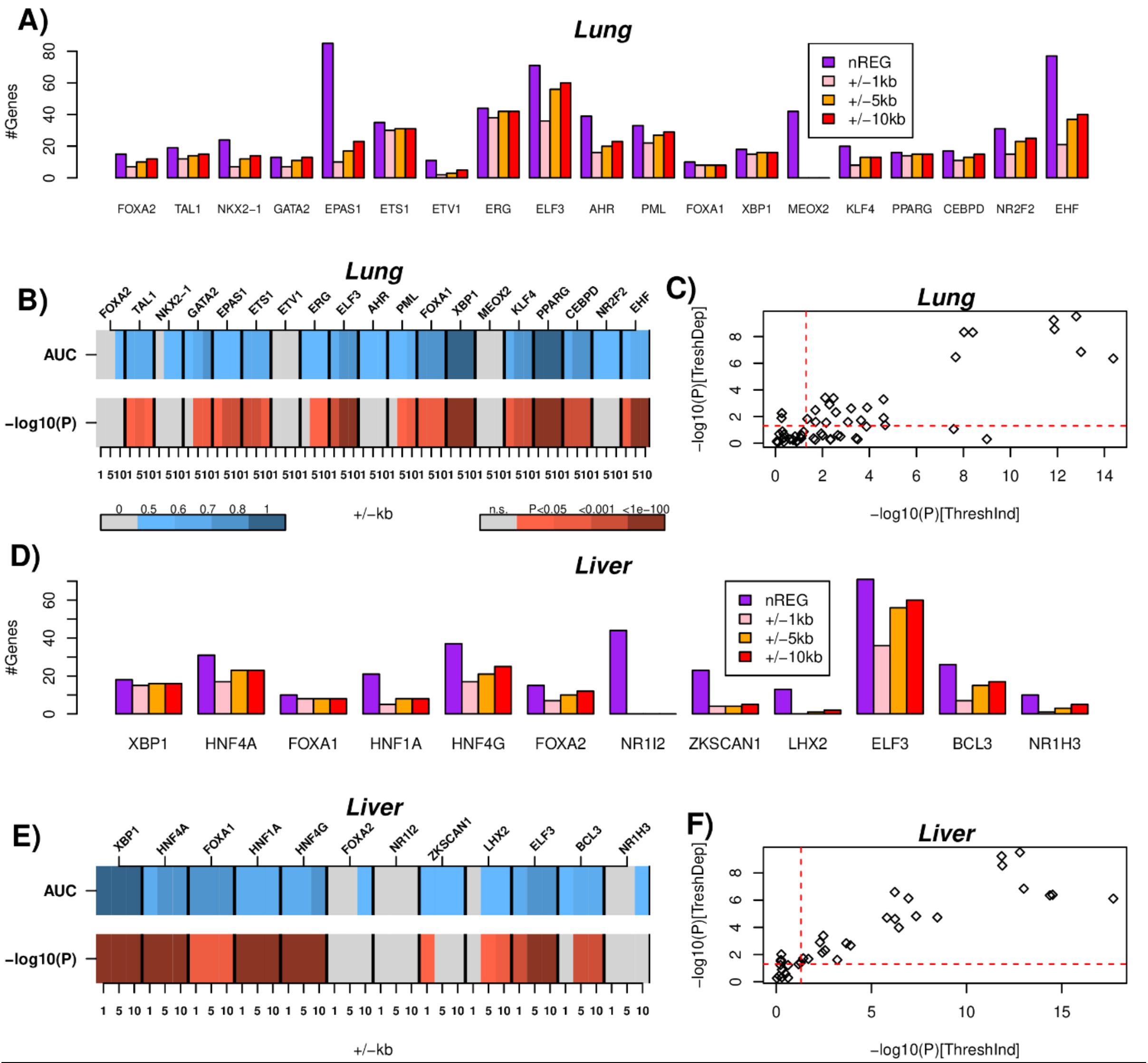


**fig.S3: Enrichment of ChIP-Seq binding targets among lung and liver regulons. A)** Barplot displaying for each of the lung-specific TFs, the number of genes in its regulon (nREG), and the number of regulon genes that are ChIP-Seq targets of the given TF within +/-1kb, +/-5kb and +/-10kb of the TSS of the gene. Only TFs for which there is available ChIP-Seq data in the ChIP-Seq atlas (http://chip-atlas.org ) were used. **B)** Threshold independent enrichment analysis using a Wilcoxon rank sum test, assessing whether the regulon-genes of a given TF have a higher ChIP-Seq binding intensity for that TF compared to genes not bound by the given TF. The Area Under the Curve (AUC) derives from the statistic of the Wilcoxon test, and the P-value is one-sided to test for overenrichment. **C)** A comparison of the –log10(P) values from the threshold-independent analysis (x-axis) vs the corresponding –log10(P) values from a threshold dependent analysis (i.e. using a threshold on the binding intensity to define ChIP-Seq targets), with the P-value derived from a binomial distribution. **D-F)** As A-C), but now for the liver-specific TFs.

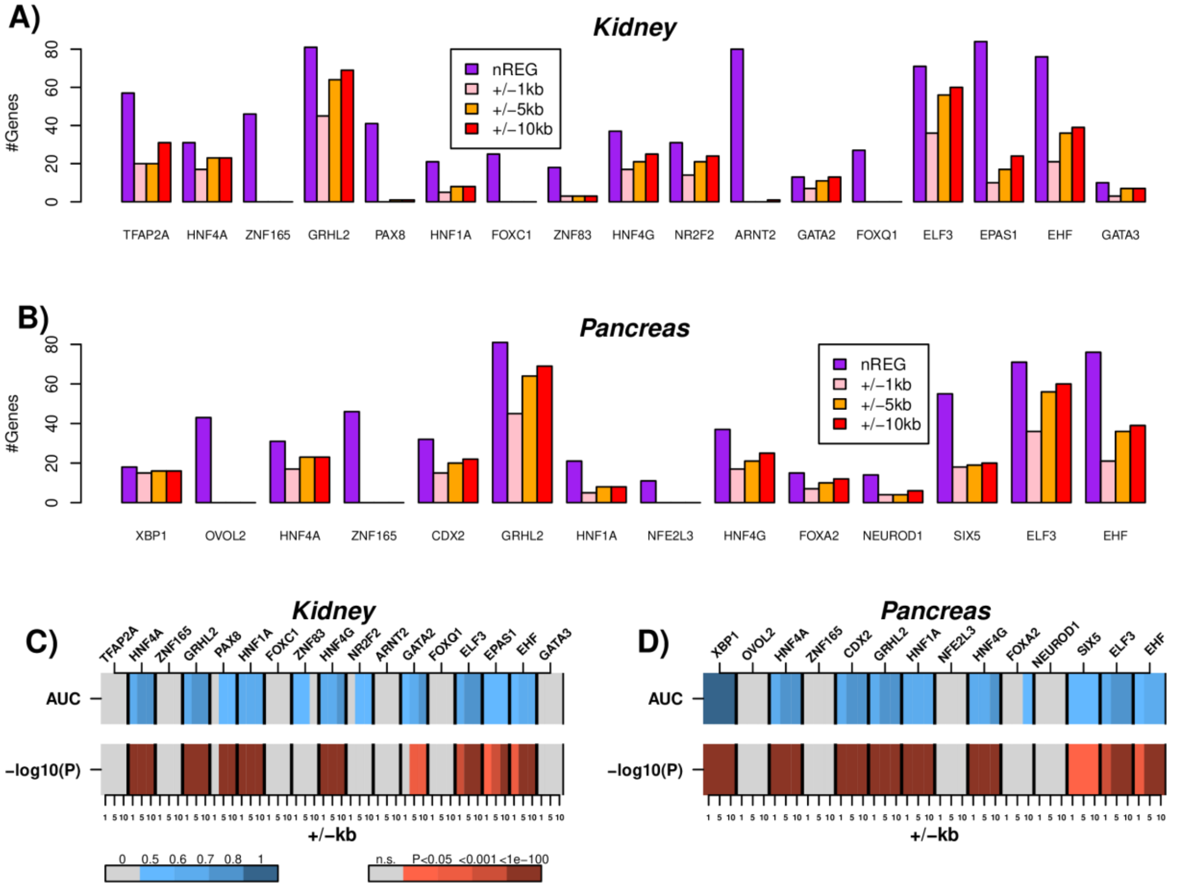


**fig.S4: Enrichment of ChIP-Seq binding targets among kidney and pancreas regulons. A)** Barplot displaying for each of the kidney-specific TFs, the number of genes in its regulon (nREG), and the number of regulon genes that are ChIP-Seq targets of the given TF within +/-1kb, +/-5kb and +/-10kb of the TSS of the gene. Only TFs for which there is available ChIP-Seq data in the ChIP-Seq atlas (http://chip-atlas.org ) were used. **B)** As A), but for pancreas. **C)** Threshold independent enrichment analysis using a Wilcoxon rank sum test, assessing whether the regulon-genes of a given kidney-specific TF have a higher ChIP-Seq binding intensity for that TF compared to genes not bound by the given TF. The Area Under the Curve (AUC) derives from the statistic of the Wilcoxon test, and the P-value is one-sided to test for overenrichment. **D)** As C), but for the pancreas-specific TFs and regulons.

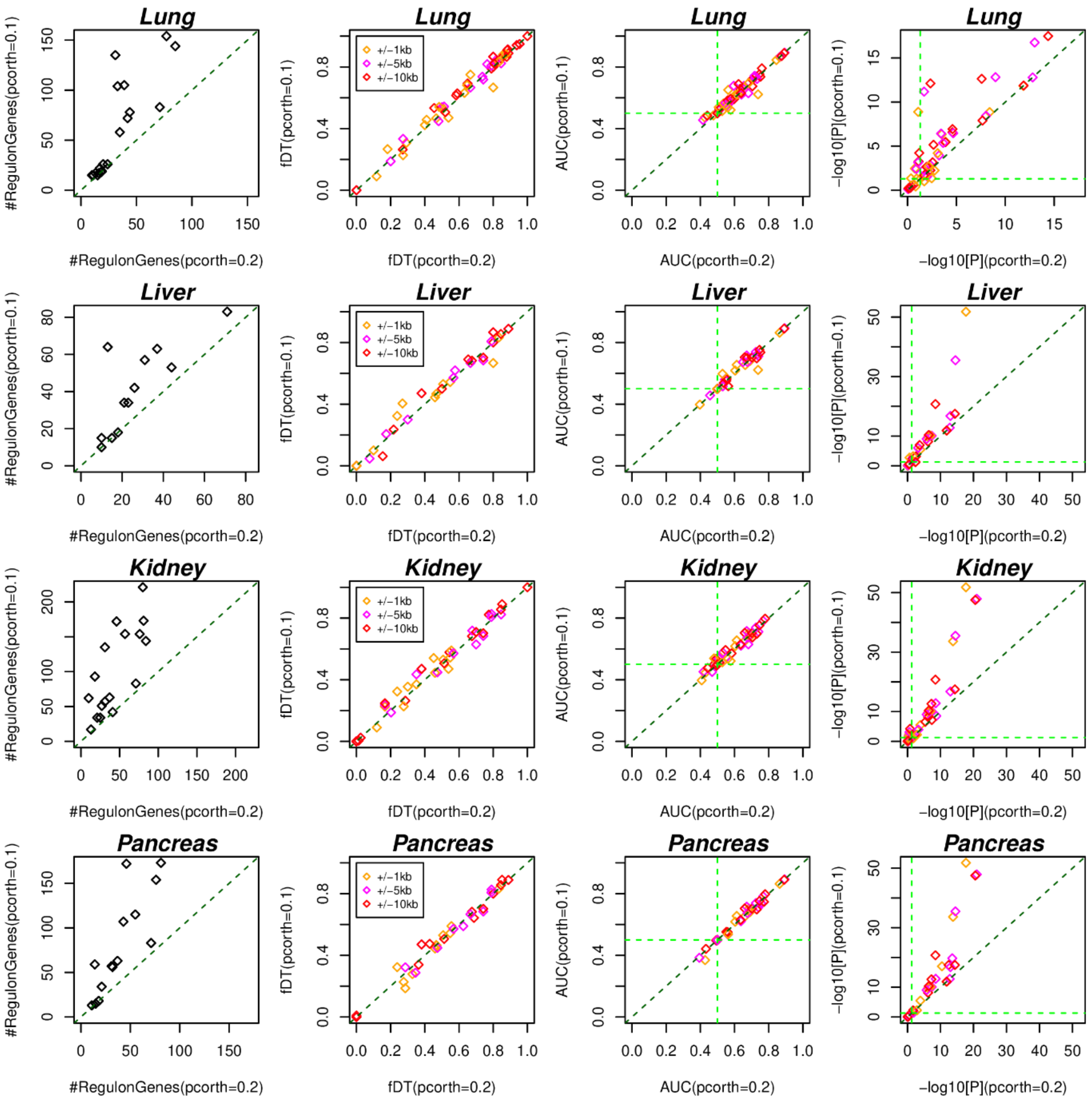


**fig.S5: Robustness of TF-binding site enrichment of SCIRA regulons to partial correlation threshold.** The panels in the first column represent scatterplots of the number of genes in the tissue-specific regulons for a partial correlation threshold of 0.2 (x-axis) vs. the corresponding number when the partial correlation threshold is set to 0.1 (y-axis). Panels in the 2$^{nd}$ column compare the fractions of regulon targets that are direct binding targets for the two different choices of partial correlation threshold. Colors label the different genomic-windows from the TSS of a regulon gene to determine whether a binding event is linked to the gene or not. Panels in the 3$^{rd}$ column compare the AUC-statistic of enrichment for TF-binding targets within the corresponding regulons for the same two partial correlation thresholds, as indicated. The AUC-statistic derives from the Wilcoxon rank sum test. Vertical and horizontal green dashed lines define the boundary of no association (AUC≤0.5). Panels in the 4$^{th}$ column compare the corresponding significance levels from the one-tailed Wilcoxon rank sum test, with the vertical and horizontal green dashed lines representing the P=0.05 significance level.

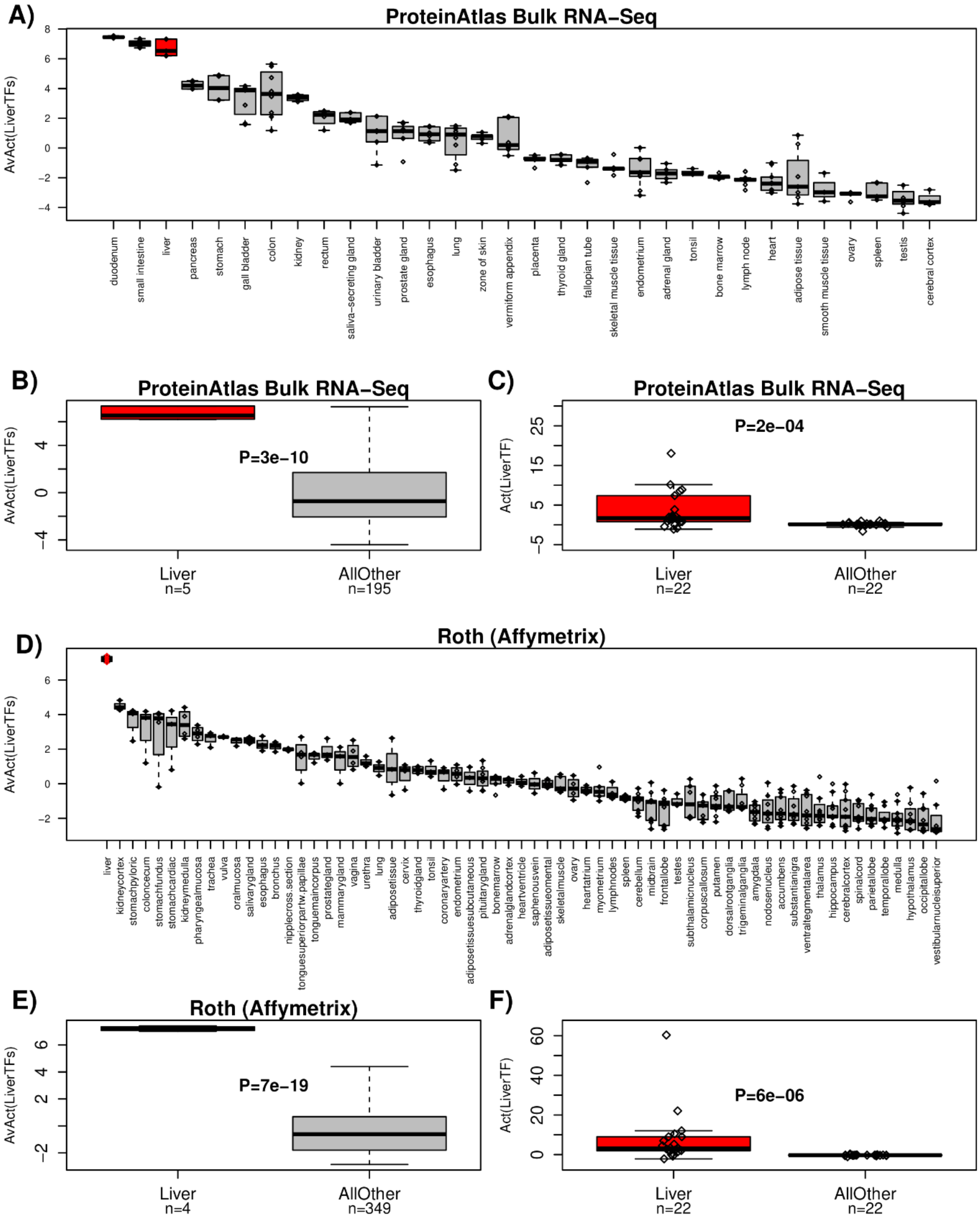


**fig.S6: Validation of the liver-specific TF regulons in independent multi-tissue expression sets. A)** Boxplots of the average estimated TF-activity level of 22 liver-specific TFs across different tissues profiled as part of the Protein Atlas project. In red we highlight the tissue "liver". **B)** Comparison boxplot of the same average activity level between liver and all other tissues. Number of tissue samples in each group is given. P-value is from a one-tailed t-test. **C)** Boxplots of the individual 22 liver-specific TF activity levels in liver vs all other tissue-types, where values across samples within a tissue have been averaged. P-value is from a one-tailed paired Wilcoxon rank sum test. **D-F)** As A-C), but now for the multi-tissue Affymetrix mRNA expression dataset from Roth et al [2].

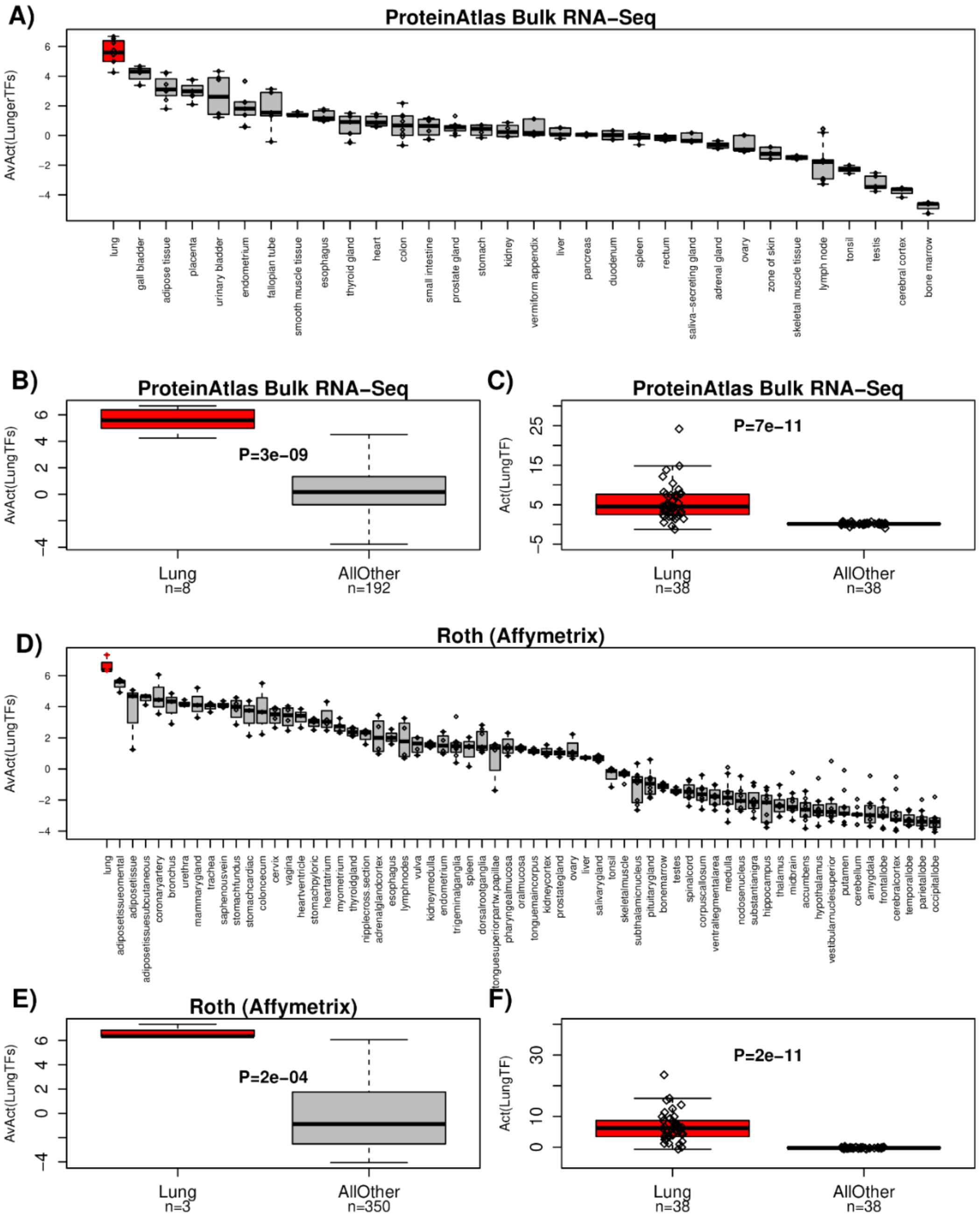


**fig.S7: Validation of the lung-specific TF regulons in independent multi-tissue expression sets. A)** Boxplots of the average estimated TF-activity level of 38 lung-specific TFs across different tissues profiled as part of the Protein Atlas project. In red we highlight the tissue "lung". **B)** Comparison boxplot of the same average activity level between lung and all other tissues. Number of tissue samples in each group is given. P-value is from a one-tailed t-test. **C)** Boxplots of the individual 38 lung-specific TF activity levels in lung vs all other tissue-types, where values across samples within a tissue have been averaged. P-value is from a one-tailed paired Wilcoxon rank sum test. **D-F)** As A-C), but now for the multi-tissue Affymetrix mRNA expression dataset from Roth et al [2].

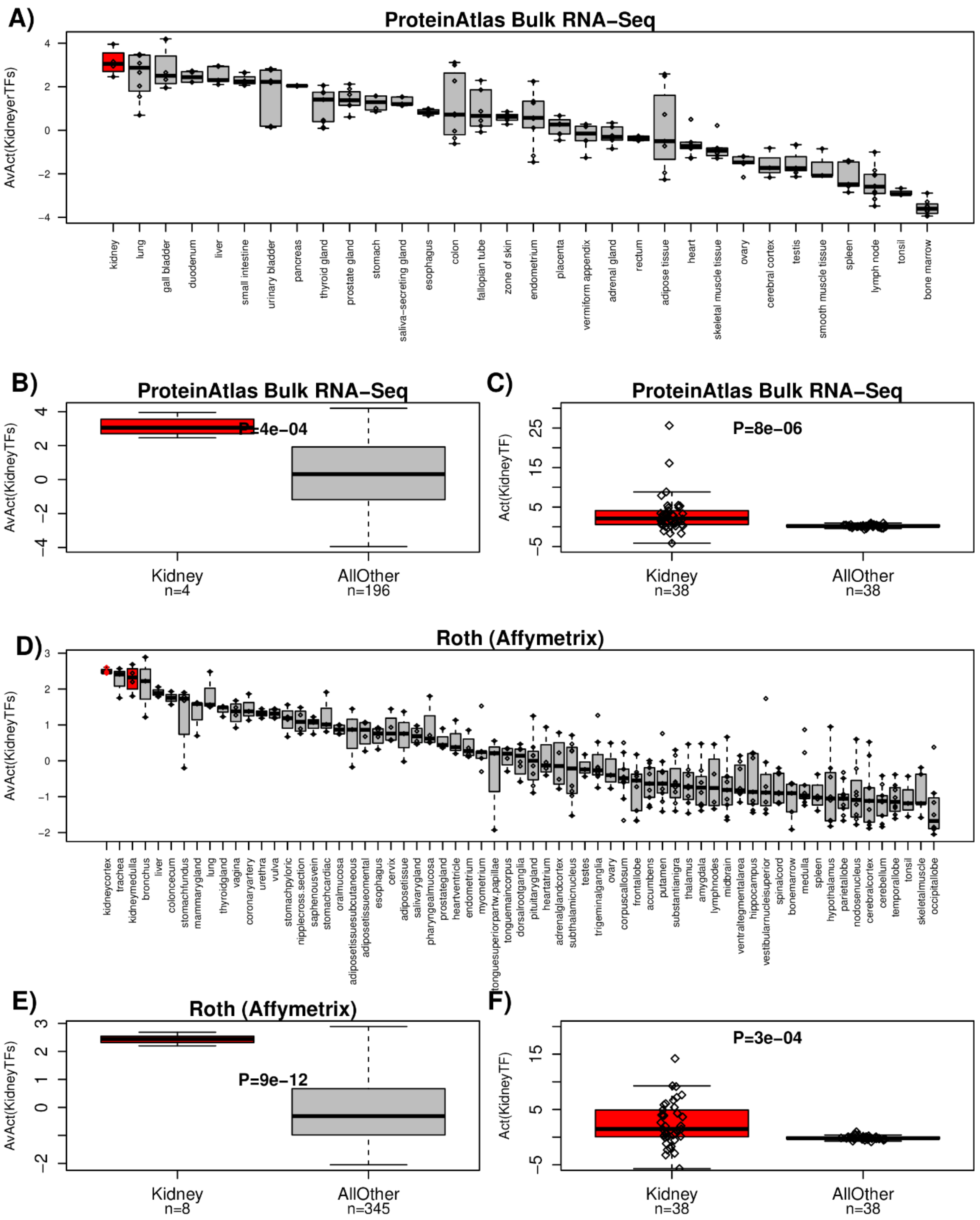


**fig.S8: Validation of the kidney-specific TF regulons in independent multi-tissue**

**expression sets. A)** Boxplots of the average estimated TF-activity level of the 38 kidney-specific TFs across different tissues profiled as part of the Protein Atlas project. In red we highlight the tissue “kidney”. **B)** Comparison boxplot of the same average activity level between kidney and all other tissues. Number of tissue samples in each group is given. P-value is from a one-tailed t-test. **C)** Boxplots of the individual 38 kidney-specific TF activity levels in kidney vs all other tissue-types, where values across samples within a tissue have been averaged. P-value is from a one-tailed paired Wilcoxon rank sum test. **D-F)** As A-C), but now for the multi-tissue Affymetrix mRNA expression dataset from Roth et al [2].

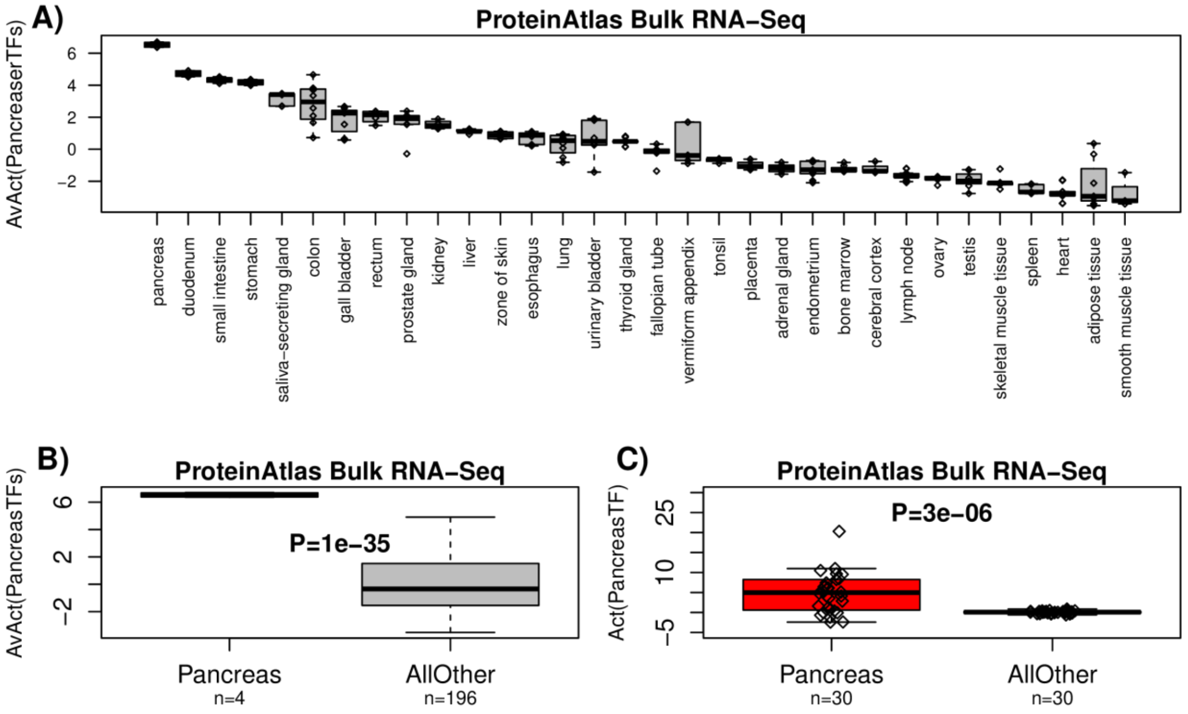


**fig.S9: Validation of the pancreas-specific TF regulons in the independent multi-tissue bulk RNA-Seq set from the ProteinAtlas. A)** Boxplots of the average estimated TF-activity level of the 30 pancreas-specific TFs across different tissues profiled as part of the Protein Atlas project. In red we highlight the tissue “pancreas”. **B)** Comparison boxplot of the same average activity level between pancreas and all other tissues. Number of tissue samples in each group is given. P-value is from a one-tailed t-test. **C)** Boxplots of the individual 30 pancreas-specific TF activity levels in pancreas vs all other tissue-types, where values across samples within a tissue have been averaged. P-value is from a one-tailed paired Wilcoxon rank sum test.

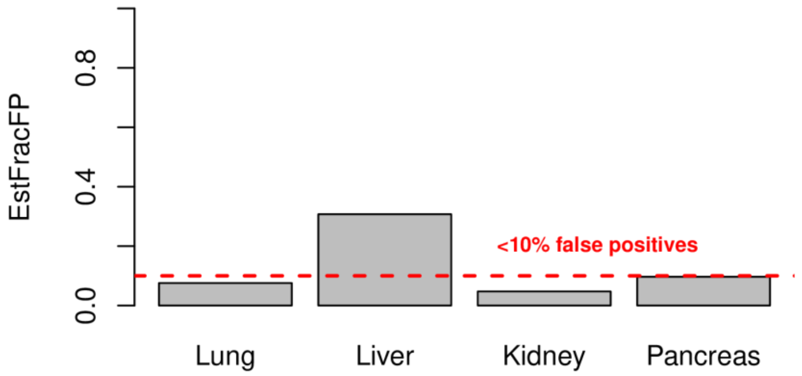


**fig.S10: Estimation of false positives in derived regulons.** Estimated fraction of false positives (EstFracFP, y-axis) within the tissue-specific regulons for 4 tissue-types (x-axis), as indicated. For a given tissue-type, the fraction of false positive regulon genes was estimated by computing differential expression statistics for all tissue-specific regulon genes in the multi-tissue bulk RNA-Seq set of the ProteinAtlas comparing the tissue of interest to all other tissue-types, and then calculating the fraction of positively correlated and negatively correlated regulon genes exhibiting downregulation and upregulation in the Protein Atlas set.

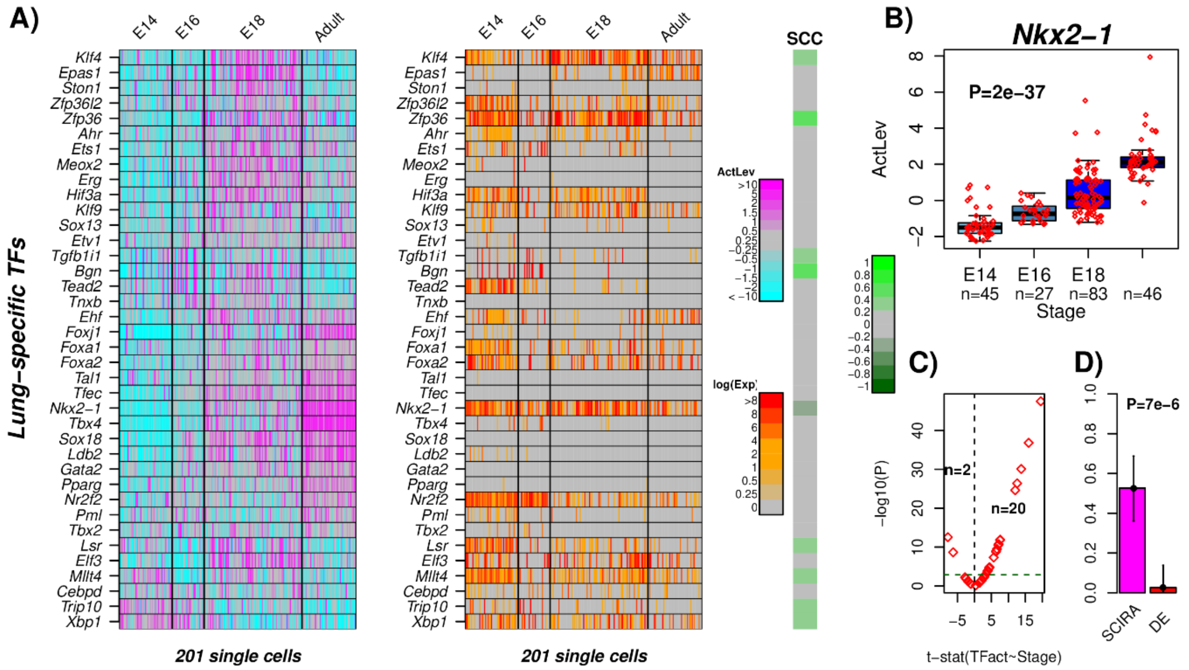


**fig.S11: Validation and improved sensitivity of SCIRA in lung development. A)** Heatmap of regulator activity of the 38 lung-specific TFs across 201 single cells representing four developmental time points in lung development in mice, as estimated using the SCIRA algorithm. Corresponding heatmap displaying $\log_2$(FPKM+1) values. Spearman Rank Correlation Coefficient (SCC) between TF-activity and TF-expression profiles. **B)** Pattern of regulatory activity change for *Nkx2-1*. P-value is from a linear regression. In each boxplot, horizontal lines describe median, interquartile range and whiskers extend to 1.5*inter-quartile range. **C)** Scatterplot of t-statistics of a linear regression of activity level against timepoint (x-axis) vs. the significance of the t-statistic [ $-\log_{10}$(P-value), y-axis] for all 38 TFs. The number of significant associations at a Bonferroni adjusted P<0.05 level are given. **D)** Barplot comparing the sensitivity (SE) to detect increased activity (SCIRA) or differential overexpression (DE) for the 38 TFs. Error bars represent 95% confidence intervals estimated using Wilson's continuity correction. P-value is from a one-tailed Fisher-exact test comparing SCIRA to DE.

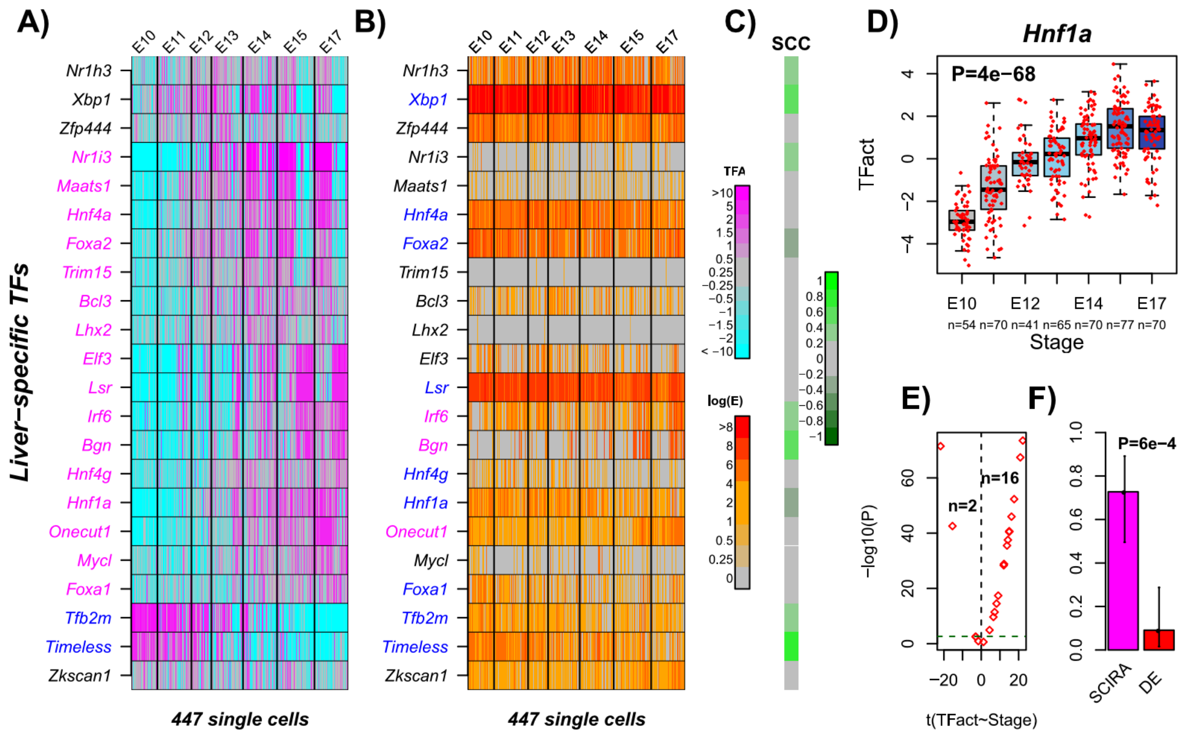


**fig.S12: Validation and improved sensitivity of SCIRA in liver development. A)** Heatmap of regulatory activity of 22 liver-specific TFs across 447 single cells representing seven developmental time points in mouse liver development (e.g E10=embryonic day 10), as estimated using the SCIRA algorithm. **B)** Corresponding heatmap displaying $\log_2$(TPM+1) values. TFs exhibiting significant increase and decrease in activity/expression are indicated in pink and blue, respectively. **C)** Color bar representing the Spearman Correlation Coefficients (SCC) between TF-activity and TF-expression. **D)** Pattern of regulatory activity change for *Hnf1a* as predicted by SCIRA. P-value is from a linear regression. In each boxplot, horizontal lines describe median, interquartile range and whiskers extend to 1.5*inter-quartile range. **E)** Scatterplot of t-statistics of a linear regression of activity level against time point (x-axis) vs. the significance of the t-statistic [ $-\log_{10}$(P-value), y-axis] for all 22 TFs. The number of significant associations at a Bonferroni adjusted P<0.05 level are given. **F)** Barplot comparing the sensitivity (SE) to detect increased activity (SCIRA) or upregulation (DE) across the 22 TFs. Error bars represent 95% confidence intervals estimated using Wilson's continuity correction. P-value is from a one-tailed Fisher-exact test comparing SCIRA to DE.

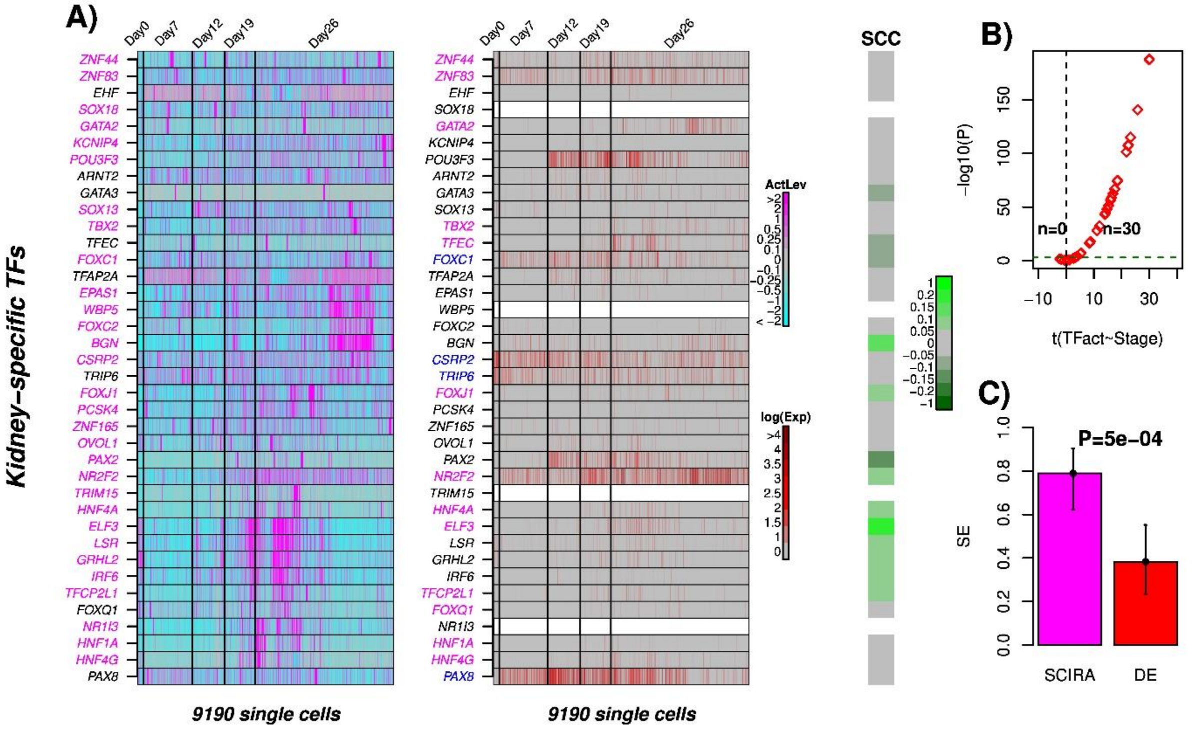


**fig.S13: Validation and improved sensitivity of SCIRA in kidney development. A)** Heatmap of regulator activity of the 38 kidney-specific TFs across 9190 single cells representing 5 differentiation stages from iPSCs (Day-0) to differentiated kidney organoid cells (Day-26), as estimated using the SCIRA algorithm. Corresponding heatmap displaying normalized expression values. Spearman Rank Correlation Coefficients (SCC) between corresponding TF-activity and TF-expression profiles. **B)** Scatterplot of t-statistics of a linear regression of activity level against timepoint (x-axis) vs. the significance of the t-statistic [ –log10(P-value), y-axis] for all 38 TFs. The number of significant associations at a Bonferroni adjusted P<0.05 level are given. **C)** Barplot comparing the sensitivity (SE) to detect increased activity (SCIRA) or upregulation (DE) for the 26 TFs with mouse homologs. Error bars represent 95% confidence intervals estimated using Wilson's continuity correction. P-value is from a one-tailed Fisher-exact test comparing SCIRA to DE.

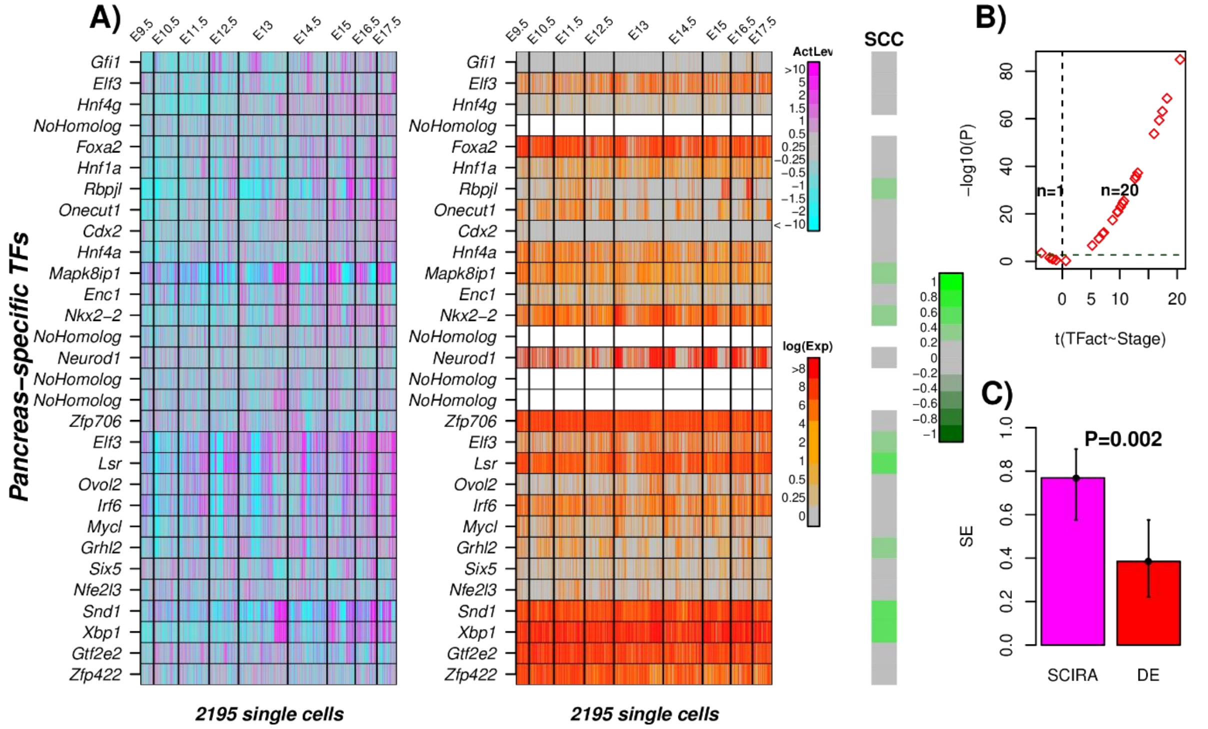


**fig.S14: Validation and improved sensitivity of SCIRA in pancreas development. A)** Heatmap of regulator activity of the 30 pancreas-specific TFs across 2195 single cells representing 9 developmental time points in pancreas development in mice, as estimated using the SCIRA algorithm. Corresponding heatmap displaying $\log_2$(TPM+1) expression values. Spearman Rank Correlation Coefficients (SCC) between corresponding TF-activity and TF-expression profiles. **B)** Scatterplot of t-statistics of a linear regression of activity level against timepoint (x-axis) vs. the significance of the t-statistic [ $-\log_{10}$(P-value), y-axis] for all 30 TFs. The number of significant associations at a Bonferroni adjusted $P<0.05$ level are given. **C)** Barplot comparing the sensitivity (SE) to detect increased activity (SCIRA) or upregulation (DE) for the 26 TFs with mouse homologs. Error bars represent 95% confidence intervals estimated using Wilson's continuity correction. P-value is from a one-tailed Fisher-exact test comparing SCIRA to DE.

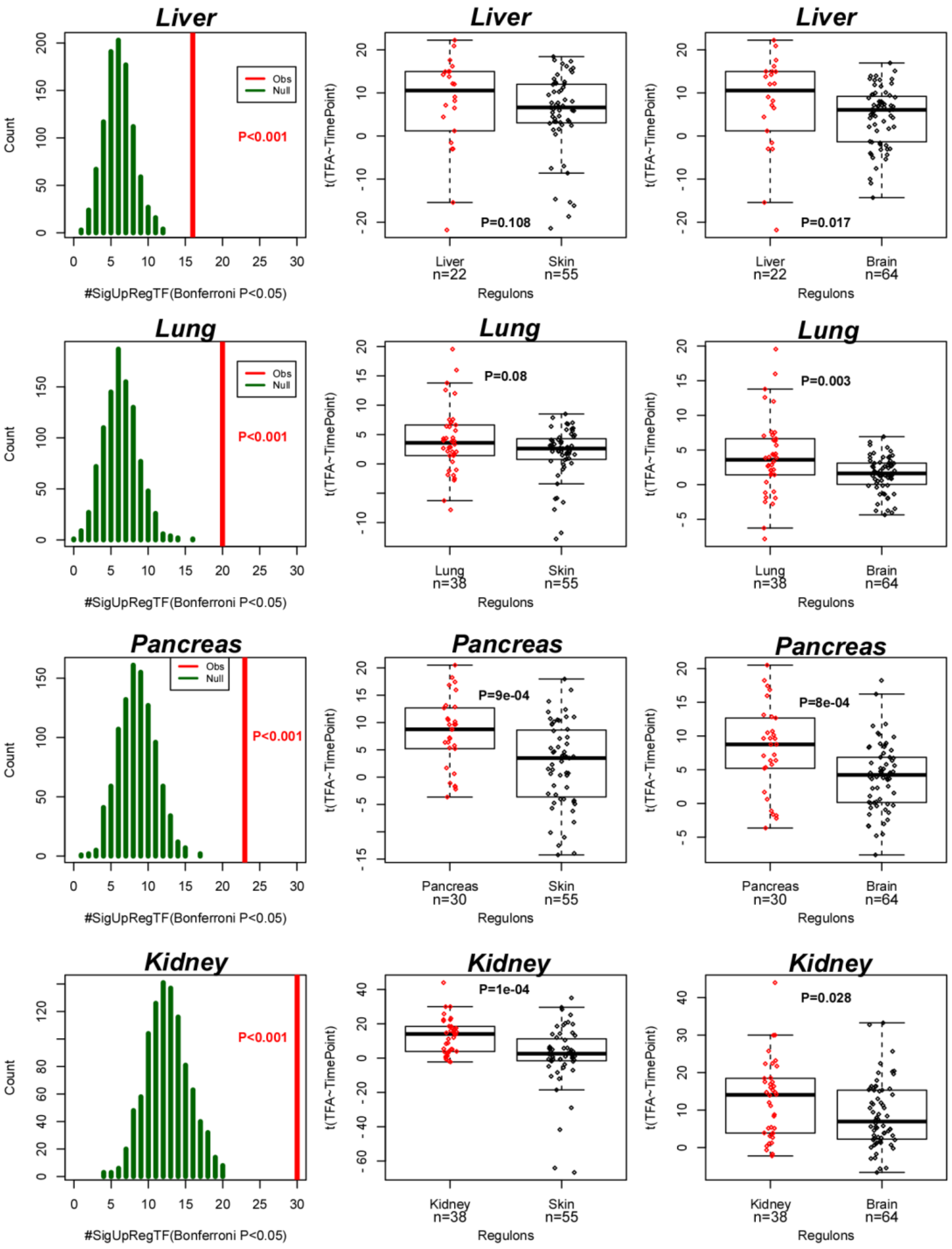


**fig.S15: Monte-Carlo Randomization and specificity of regulons.** Left panels: for each tissue-type we compare the number of tissue-specific TFs predicted by SCIRA to be upregulated during development/differentiation (vertical red line) to the null distribution (1000

Monte-Carlo runs, green lines) obtained by randomly constructing regulons of the same size and same distribution of positive and negative interactions. In the plot for pancreas all TF-regulons were considered, not just those with mouse homologs. Middle and right panels compare the t-statistics of association of TF-activity (as estimated using SCIRA regulons) with developmental stage/timepoint for the tissue-specific regulons to skin and brain specific regulons. P-value is from a one-tailed Wilcoxon rank sum test.

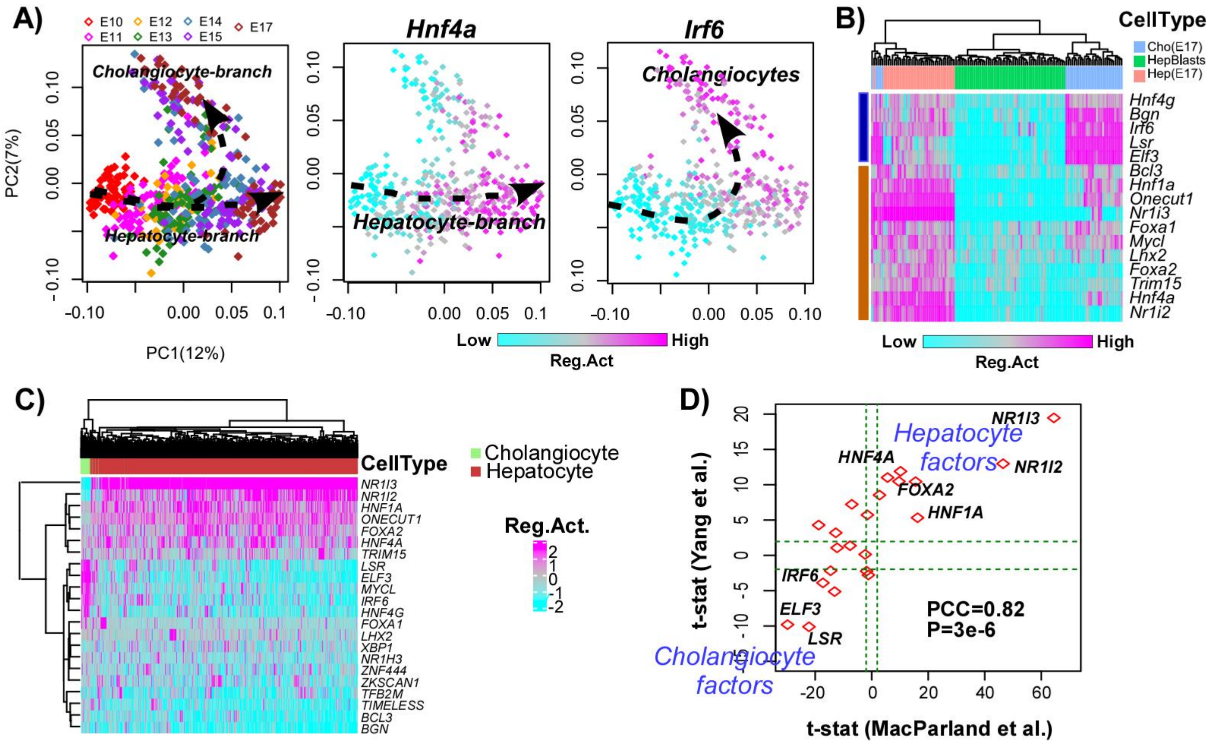


**fig.S16: SCIRA exhibits power to detect TFs of minor cell-types and demonstrates cell-type specificity. A)** PCA scatterplot of the 447 single cells derived from a timecourse of hepatoblasts into hepatocytes and cholangiocytes, as obtained by applying PCA on the regulatory activity matrix shown in Fig.2A. From left to right, cells are labeled according to developmental stage (embryonic-days E10 to E17), regulatory activity of *Hnf4a* and *Irf6*. **B)** Clustering heatmap over the 16 TFs exhibiting increased activity during differentiation, and over all single cells at the start (E10) and endpoints (E17). Cell-types annotated as hepatoblasts (E10, Hepblasts), hepatocytes at E17 (Hep(E17)) and cholangiocytes (Cho(E17)). **C)** Hierarchical clustering of the 22 liver-specific TFs and single cells over the regulatory activity matrix as estimated by SCIRA in the 10X scRNA-Seq dataset of MacParland. Two main cell clusters are annotated by cell-type. **D)** Scatterplot of t-statistics of differential regulatory activity between hepatoblasts and cholangiocytes as calculated in Yang et al differentiation timecourse scRNA-Seq set (y-axis) vs. the ones calculated in the 10X MacParland et al set (x-axis). Pearson Correlation Coefficient (PCC) and P-value are given.

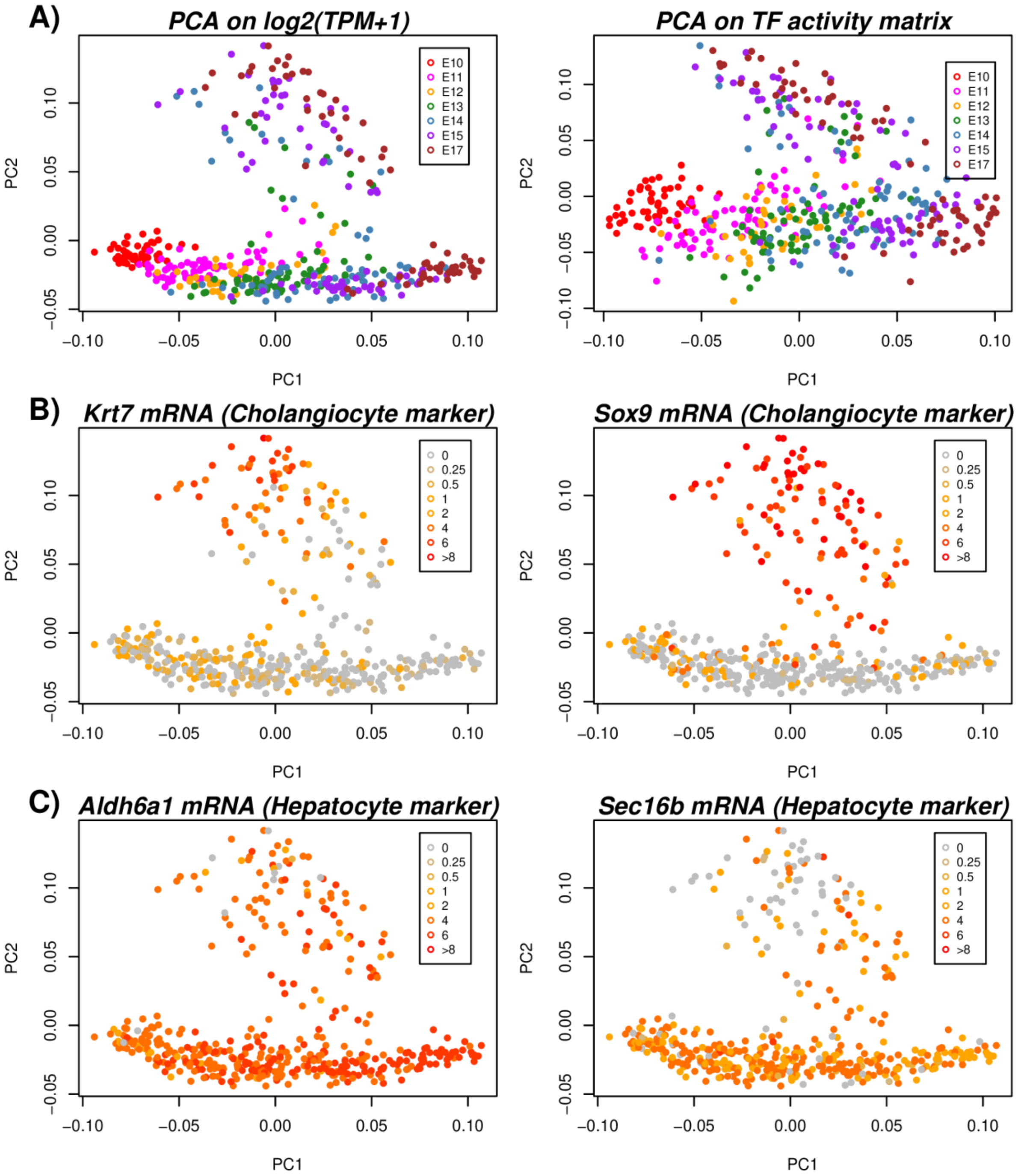


**fig.S17: PCA analysis on liver scRNA-Seq set and definition of cholangiocyte and hepatocyte branches. A)** Left panel depicts the PCA scatterplot of a PCA on the log2(TPM+1) expression matrix over all genes of the Yang et al liver scRNA-Seq study [3]. Right panel is the corresponding PCA scatterplot of a PCA on the transcription factor activity matrix over the 22 liver-specific TFs as estimated using SCIRA, which largely recapitulates the pattern derived from the full expression matrix. **B)** PCA scatterplot as in left panel of A), but now with cells colored according to the level of expression of two cholangiocyte markers (*Krt7, Sox9*) as identified in MacParland et al [4], defining the cholangiocyte branch. **C)** As B), but now for two hepatocyte markers (*Aldh6a1, Sec16b*), as identified in MacParland et al.

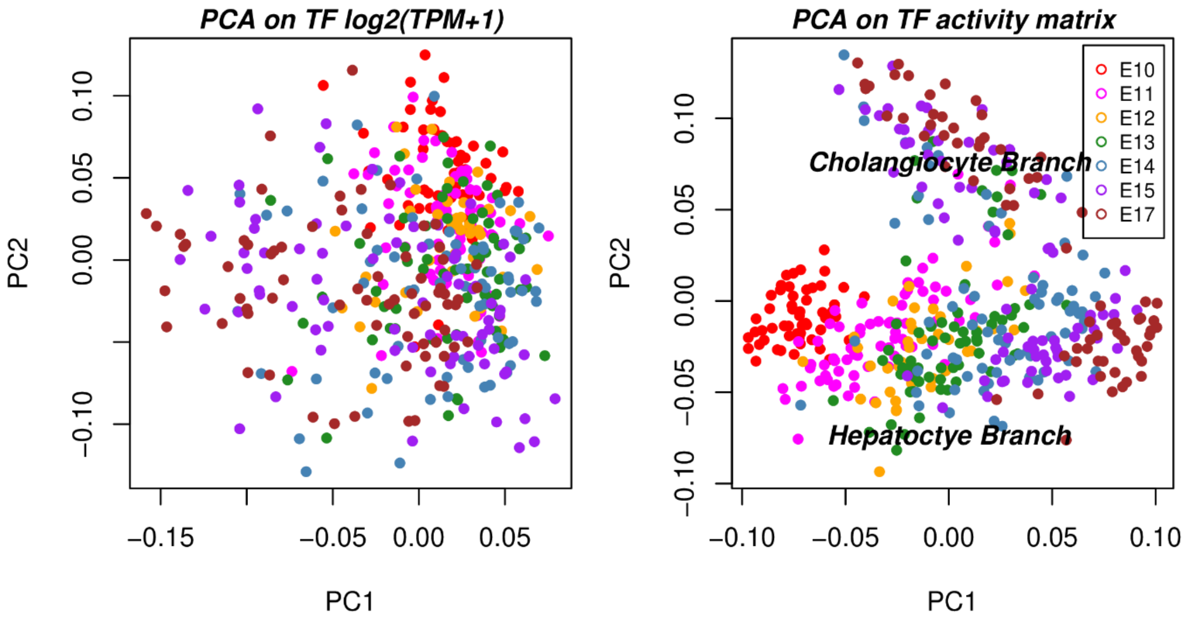


**fig.S18: Comparative PCA analysis of TF expression and TF activity matrices.** Left panel depicts the PCA scatterplot of a PCA on the 22 TF times 447 single-cell log2(TPM+1) expression matrix of the Yang et al liver scRNA-Seq study [3]. Right panel is the corresponding PCA scatterplot of a PCA on the transcription factor activity matrix over the 22 liver-specific TFs as estimated using SCIRA.

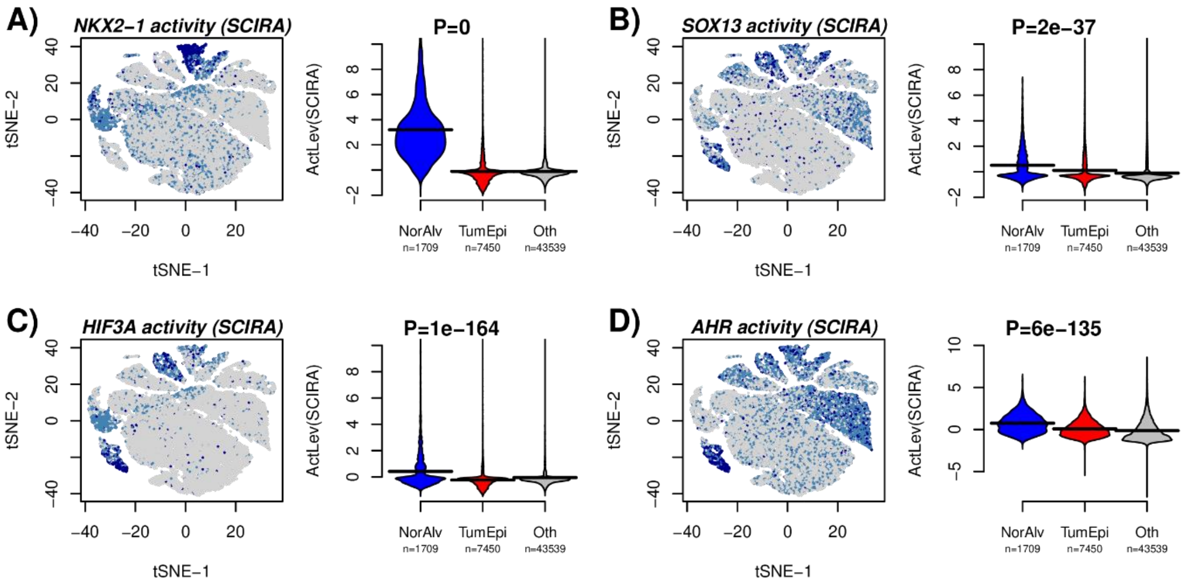


**fig.S19: Inactivation of lung-specific TFs in lung tumor epithelial cells. A)** t-SNE scatterplot of approximately 52,000 single cells from 5 lung cancer patients, with cells color-labeled according to the SCIRA predicted activity of *NKX2-1*. Right panel shows beanplots of the predicted SCIRA activity level of *NKX2-1* between normal alveolar, tumor epithelial and all other cells. P-value is from a linear model with activity level as response and normal alveolar/tumor epithelial status as predictor. P=0 means P<1e-500. **B-D)** As A), but now for the other lung-specific TFs *SOX13, HIF3A* and *AHR*.

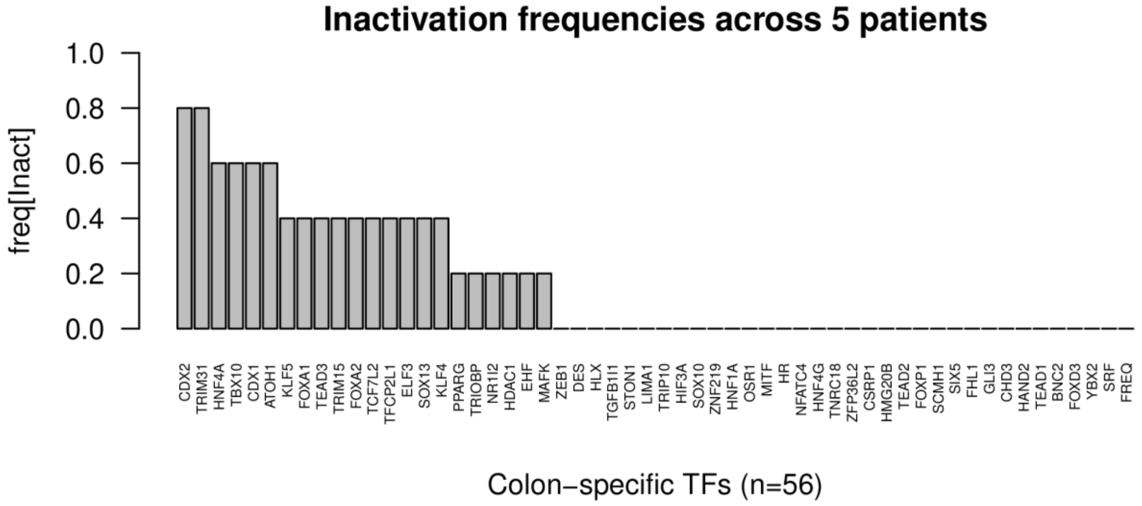


**fig.S20: Frequency of inactivation of colon-specific TFs across 5 patients.** Barplots displaying the frequency of inactivation of all 56 colon specific TFs, ranked by their frequency. Inactivation was determined for each of the 5 patients separately, by comparing the SCIRA TF-activity estimates of the cancer cells to those of the normal cells within the patient. Bonferroni-adjusted $P < 0.05$ on the t-test comparing these activity estimates was used to declare inactivation.

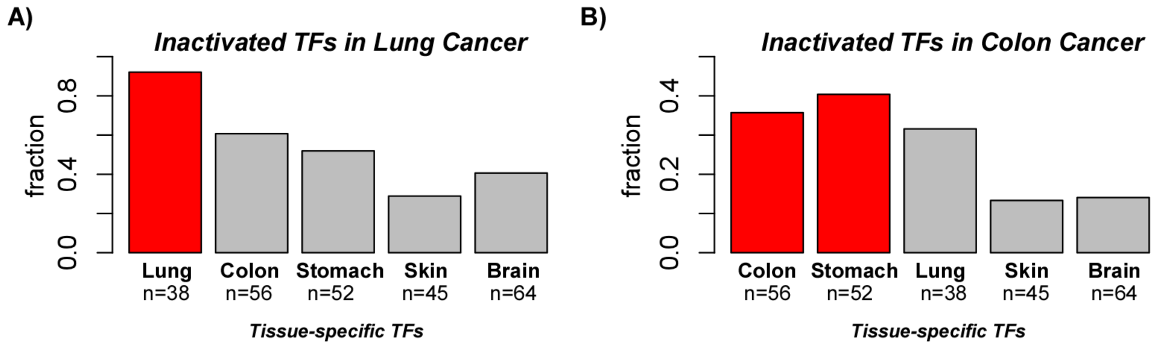


**fig.S21: Tissue-specificity of TF-inactivation in cancer cells. A)** Barplot displaying the fraction of tissue-specific TFs that are significant inactivated (Bonferroni adjusted $P<0.05$) in the single lung cancer cells compared to single normal cells. **B)** As A), but now in the scRNA-Seq dataset of normal and cancer colon cells. The number of tissue-specific TFs is given below bars.

### SUPPLEMENTARY TABLES

| TF(Human) | TF(Mouse) | # in Regulon | Positive | Inhibitory |
|---|---|---|---|---|
| *XBP1* | *Xbp1* | 18 | 18 | 0 |
| *LSR* | *Lsr* | 72 | 72 | 0 |
| *HNF4A* | *Hnf4a* | 31 | 28 | 3 |
| *BGN* | *Bgn* | 93 | 93 | 0 |
| *FOXA1* | *Foxa1* | 10 | 10 | 0 |
| *ONECUT1* | *Onecut1* | 15 | 15 | 0 |
| *HNF1A* | *Hnf1a* | 21 | 21 | 0 |
| *IRF6* | *Irf6* | 53 | 53 | 0 |
| *TIMELESS* | *Timeless* | 29 | 29 | 0 |
| *MYCL1* | *Mycl* | 40 | 38 | 2 |
| *TRIM15* | *Trim15* | 12 | 11 | 1 |
| *HNF4G* | *Hnf4g* | 37 | 28 | 9 |
| *FOXA2* | *Foxa2* | 15 | 15 | 0 |
| *NR1I2* | *Nr1i2* | 44 | 42 | 2 |
| *NR1I3* | *Nr1i3* | 151 | 151 | 0 |
| *ZKSCAN1* | *Zkscan1* | 23 | 23 | 0 |
| *TFB2M* | *Tfb2m* | 101 | 101 | 0 |
| *LHX2* | *Lhx2* | 13 | 12 | 1 |
| *ELF3* | *Elf3* | 71 | 71 | 0 |
| *BCL3* | *Bcl3* | 26 | 26 | 0 |
| *ZNF444* | *Zfp444* | 23 | 23 | 0 |
| *NR1H3* | *Nr1h3* | 10 | 10 | 0 |

**table.S1: Summary of the liver-specific regulatory network.** Table lists the human gene symbol and mouse homolog of the 22 liver-specific transcription factors (TFs) for the liver-specific regulatory network derived using the SEPIRA algorithm [5] on the GTEX dataset [6]. The other columns list the number of gene targets in the TF-regulon, and the number of these that represent positive and inhibitory interactions.

| TF(Human) | TF(Mouse) | # in Regulon | Positive | Inhibitory |
|---|---|---|---|---|
| *TFEC* | *Tfec* | 33 | 33 | 0 |
| *TBX2* | *Tbx2* | 18 | 14 | 4 |
| *FOXA2* | *Foxa2* | 15 | 15 | 0 |
| *TAL1* | *Tal1* | 19 | 19 | 0 |
| *TBX4* | *Tbx4* | 16 | 16 | 0 |
| *NKX2-1* | *Nkx2-1* | 24 | 24 | 0 |
| *GATA2* | *Gata2* | 13 | 13 | 0 |
| *EPAS1* | *Epas1* | 85 | 83 | 2 |
| *FOXJ1* | *Foxj1* | 152 | 152 | 0 |
| *LDB2* | *Ldb2* | 63 | 63 | 0 |
| *ETS1* | *Ets1* | 35 | 35 | 0 |
| *ETV1* | *Etv1* | 11 | 11 | 0 |
| *ERG* | *Erg* | 44 | 44 | 0 |
| *ELF3* | *Elf3* | 71 | 71 | 0 |
| *SOX13* | *Sox13* | 14 | 14 | 0 |
| *AHR* | *Ahr* | 39 | 38 | 1 |
| *PML* | *Pml* | 33 | 28 | 5 |
| *FOXA1* | *Foxa1* | 10 | 10 | 0 |
| *MLLT4* | *Mllt4* | 26 | 26 | 0 |
| *BGN* | *Bgn* | 93 | 93 | 0 |
| *ZFP36* | *Zfp36* | 19 | 18 | 1 |
| *TNXB* | *Tnxb* | 40 | 40 | 0 |
| *SOX18* | *Sox18* | 60 | 60 | 0 |
| *TEAD2* | *Tead2* | 53 | 52 | 1 |
| *XBP1* | *Xbp1* | 18 | 18 | 0 |
| *MEOX2* | *Meox2* | 42 | 41 | 1 |
| *KLF4* | *Klf4* | 20 | 20 | 0 |
| *HIF3A* | *Hif3a* | 10 | 10 | 0 |
| *LSR* | *Lsr* | 70 | 70 | 0 |
| *KLF9* | *Klf9* | 15 | 15 | 0 |
| *STON1* | *Ston1* | 31 | 30 | 1 |
| *PPARG* | *Pparg* | 16 | 16 | 0 |
| *ZFP36L2* | *Zfp36l2* | 24 | 24 | 0 |
| *CEBPD* | *Cebpd* | 17 | 17 | 0 |
| *TRIP10* | *Trip10* | 42 | 23 | 19 |
| *NR2F2* | *Nr2f2* | 31 | 24 | 7 |
| *TGFB1I1* | *Tgfb1i1* | 112 | 108 | 4 |
| *EHF* | *Ehf* | 77 | 50 | 27 |

**table.S2: Summary of the lung-specific regulatory network.** Table lists the human gene symbol and mouse homolog of the 38 lung-specific transcription factors (TFs) for the lung-specific regulatory network derived using the SEPIRA algorithm [5] on the GTEX dataset [6]. The other columns list the number of gene targets in the TF-regulon, and the number of these that represent positive and inhibitory interactions.

| TF(Human) | EntrezID | # in Regulon | Positive | Inhibitory |
|---|---|---|---|---|
| *TFAP2A* | 7020 | 57 | 9 | 48 |
| *LSR* | 51599 | 72 | 72 | 0 |
| *HNF4A* | 3172 | 31 | 28 | 3 |
| *TFEC* | 22797 | 33 | 33 | 0 |
| *ZNF165* | 7718 | 46 | 46 | 0 |
| *BGN* | 633 | 93 | 93 | 0 |
| *TBX2* | 6909 | 20 | 15 | 5 |
| *GRHL2* | 79977 | 81 | 60 | 21 |
| *PAX8* | 7849 | 41 | 39 | 2 |
| *HNF1A* | 6927 | 21 | 21 | 0 |
| *FOXC1* | 2296 | 25 | 25 | 0 |
| *CSRP2* | 1466 | 19 | 19 | 0 |
| *IRF6* | 3664 | 53 | 53 | 0 |
| *ZNF83* | 55769 | 18 | 18 | 0 |
| *ZNF44* | 51710 | 10 | 10 | 0 |
| *TRIM15* | 89870 | 12 | 11 | 1 |
| *OVOL1* | 5017 | 68 | 64 | 4 |
| *HNF4G* | 3174 | 37 | 28 | 9 |
| *NR2F2* | 7026 | 31 | 22 | 9 |
| *WBP5* | 51186 | 84 | 84 | 0 |
| *ARNT2* | 9915 | 80 | 64 | 16 |
| *NR1I3* | 9970 | 151 | 151 | 0 |
| *TFCP2L1* | 29842 | 26 | 26 | 0 |
| *GATA2* | 2624 | 13 | 13 | 0 |
| *FOXQ1* | 94234 | 27 | 27 | 0 |
| *PAX2* | 5076 | 10 | 10 | 0 |
| *ELF3* | 1999 | 71 | 71 | 0 |
| *EPAS1* | 2034 | 84 | 82 | 2 |
| *EHF* | 26298 | 76 | 49 | 27 |
| *SOX13* | 9580 | 14 | 14 | 0 |
| *TRIP6* | 7205 | 53 | 49 | 4 |
| *POU3F3* | 5455 | 19 | 15 | 4 |
| *PCSK4* | 54760 | 197 | 184 | 13 |
| *SOX18* | 54345 | 59 | 59 | 0 |
| *KCNIP4* | 80333 | 105 | 98 | 7 |
| *GATA3* | 2625 | 10 | 10 | 0 |
| *FOXJ1* | 2302 | 152 | 152 | 0 |
| *FOXC2* | 2303 | 26 | 26 | 0 |

**table.S3: Summary of the kidney-specific regulatory network.** Table lists the human gene symbol and Entrez gene ID of the 38 kidney-specific transcription factors (TFs) in the kidney-specific regulatory network derived using the SEPIRA algorithm [5] on the GTEX dataset [6]. The other columns list the number of gene targets in the TF-regulon, and the number of these that represent positive and inhibitory interactions.

| TF(Human) | TF(Mouse) | # in Regulon | Positive | Inhibitory |
|---|---|---|---|---|
| *XBP1* | *Xbp1* | 18 | 18 | 0 |
| *OVOL2* | *Ovol2* | 43 | 41 | 2 |
| *LSR* | *Lsr* | 72 | 72 | 0 |
| *HNF4A* | *Hnf4a* | 31 | 28 | 3 |
| *ZNF165* | *NA* | 46 | 46 | 0 |
| *CDX2* | *Cdx2* | 32 | 24 | 8 |
| *ZNF432* | *NA* | 18 | 18 | 0 |
| *GRHL2* | *Grhl2* | 81 | 60 | 21 |
| *ONECUT1* | *Onecut1* | 15 | 15 | 0 |
| *HNF1A* | *Hnf1a* | 21 | 21 | 0 |
| *IRF6* | *Irf6* | 53 | 53 | 0 |
| *NFE2L3* | *Nfe2l3* | 11 | 11 | 0 |
| *MYCL1* | *Mycl* | 40 | 38 | 2 |
| *ZNF22* | *Zfp422* | 12 | 12 | 0 |
| *HNF4G* | *Hnf4g* | 37 | 28 | 9 |
| *FOXA2* | *Foxa2* | 15 | 15 | 0 |
| *NKX2-2* | *Nkx2-2* | 23 | 17 | 6 |
| *GTF2E2* | *Gtf2e2* | 50 | 50 | 0 |
| *ENC1* | *Enc1* | 27 | 27 | 0 |
| *RBPJL* | *Rbpjl* | 59 | 59 | 0 |
| *NEUROD1* | *Neurod1* | 14 | 14 | 0 |
| *SIX5* | *Six5* | 55 | 51 | 4 |
| *ELF3* | *Elf3* | 71 | 71 | 0 |
| *MAPK8IP1* | *Mapk8ip1* | 171 | 169 | 2 |
| *EHF* | *Elf3* | 76 | 49 | 27 |
| *ZNF85* | *NA* | 21 | 21 | 0 |
| *SND1* | *Snd1* | 114 | 114 | 0 |
| *ZNF33B* | *NA* | 19 | 19 | 0 |
| *GFI1* | *Gfi1* | 14 | 14 | 0 |
| *ZNF706* | *Zfp706* | 21 | 21 | 0 |

**table.S4: Summary of the pancreas-specific regulatory network.** Table lists the human gene symbol and mouse homolog of the 30 pancreas-specific transcription factors (TFs) in the pancreas-specific regulatory network derived using the SEPIRA algorithm [5] on the GTEX dataset [6]. The other columns list the number of gene targets in the TF-regulon, and the number of these that represent positive and inhibitory interactions.

| Study | Species | Technology | Tissue | # Cells | # Stages/Timepoints |
|---|---|---|---|---|---|
| Treutlein et al | Mouse | Fluidigm C1 | Lung | 201 | 4 (E14 -> Adult) |
| Yang et al | Mouse | Fluidigm C1 | Liver | 447 | 7 (E10 -> E17) |
| Wu et al | Human | DropSeq | Kidney (organoid) | 9190 | 5 (Day0 -> Day26) |
| Yu et al | Mouse | Smart-Seq2 | Pancreas | 2195 | 9 (E9.5 -> E17.5) |

**table.S5: Summary of the scRNA-Seq time course differentiation/development studies analysed in this work.** Table lists the name of the study, the species, the scRNA-Seq technology, the tissue-type, the number of cells used after quality control, and the number of developmental stages/timepoints considered.

| TF(Human) | EntrezID | # in Regulon | Positive | Inhibitory |
|---|---|---|---|---|
| *ZEB1* | 6935 | 37 | 37 | 0 |
| *DES* | 1674 | 38 | 38 | 0 |
| *HLX* | 3142 | 11 | 11 | 0 |
| *PPARG* | 5468 | 17 | 17 | 0 |
| *TGFB1I1* | 7041 | 112 | 106 | 6 |
| *STON1* | 11037 | 32 | 31 | 1 |
| *LIMA1* | 51474 | 57 | 57 | 0 |
| *KLF5* | 688 | 41 | 23 | 18 |
| *HNF4A* | 3172 | 31 | 28 | 3 |
| *TRIP10* | 9322 | 41 | 22 | 19 |
| *CDX2* | 1045 | 32 | 24 | 8 |
| *HIF3A* | 64344 | 10 | 10 | 0 |
| *TRIOBP* | 11078 | 28 | 26 | 2 |
| *SOX10* | 6663 | 36 | 36 | 0 |
| *FOXA1* | 3169 | 10 | 10 | 0 |
| *ZNF219* | 51222 | 14 | 14 | 0 |
| *HNF1A* | 6927 | 21 | 21 | 0 |
| *TBX10* | 347853 | 23 | 21 | 2 |
| *TEAD3* | 7005 | 51 | 41 | 10 |
| *OSR1* | 130497 | 20 | 20 | 0 |
| *TRIM15* | 89870 | 12 | 11 | 1 |
| *MITF* | 4286 | 11 | 11 | 0 |
| *HR* | 55806 | 22 | 22 | 0 |
| *NFATC4* | 4776 | 32 | 17 | 15 |

| | | | | |
|---|---|---|---|---|
| *HNF4G* | 3174 | 37 | 28 | 9 |
| *CDX1* | 1044 | 18 | 16 | 2 |
| *TNRC18* | 84629 | 27 | 27 | 0 |
| *ZFP36L2* | 678 | 26 | 26 | 0 |
| *CSRP1* | 1465 | 89 | 89 | 0 |
| *FOXA2* | 3170 | 15 | 15 | 0 |
| *HMG20B* | 10362 | 20 | 15 | 5 |
| *TCF7L2* | 6934 | 44 | 43 | 1 |
| *TEAD2* | 8463 | 55 | 53 | 2 |
| *FOXP1* | 27086 | 10 | 10 | 0 |
| *NR1I2* | 8856 | 44 | 42 | 2 |
| *HDAC1* | 3065 | 34 | 31 | 3 |
| *TFCP2L1* | 29842 | 26 | 26 | 0 |
| *SCMH1* | 22955 | 33 | 30 | 3 |
| *SIX5* | 147912 | 55 | 51 | 4 |
| *FHL1* | 2273 | 64 | 64 | 0 |
| *ELF3* | 1999 | 71 | 71 | 0 |
| *GLI3* | 2737 | 33 | 33 | 0 |
| *CHD3* | 1107 | 14 | 13 | 1 |
| *HAND2* | 9464 | 15 | 15 | 0 |
| *EHF* | 26298 | 76 | 49 | 27 |
| *MAFK* | 7975 | 10 | 10 | 0 |
| *TEAD1* | 7003 | 61 | 60 | 1 |
| *TRIM31* | 11074 | 43 | 42 | 1 |
| *ATOH1* | 474 | 35 | 32 | 3 |
| *BNC2* | 54796 | 21 | 21 | 0 |
| *FOXD3* | 27022 | 29 | 29 | 0 |
| *YBX2* | 51087 | 32 | 31 | 1 |
| *SOX13* | 9580 | 14 | 14 | 0 |
| *SRF* | 6722 | 45 | 45 | 0 |
| *FREQ* | 23413 | 56 | 56 | 0 |
| *KLF4* | 9314 | 19 | 19 | 0 |

**table.S6: Summary of the colon-specific regulatory network.** Table lists the human gene symbol and Entrez gene ID of the 56 colon-specific transcription factors (TFs) in the colon-specific regulatory network derived using the SEPIRA algorithm [5] on the GTEX dataset [6].
